# Bidirectional regulation of cognitive and anxiety-like behaviors by dentate gyrus mossy cells in male and female mice

**DOI:** 10.1101/2020.07.05.188664

**Authors:** Justin J Botterill, K Yaragudri Vinod, Kathleen J Gerencer, Cátia M Teixeira, John J LaFrancois, Helen E Scharfman

## Abstract

The dentate gyrus (DG) of the hippocampus is important for cognitive and affective behaviors. However, the circuits underlying these behaviors are unclear. DG mossy cells (MCs) have been a focus of attention because of their excitatory synapses on the primary DG cell type, granule cells (GCs). However, MCs also activate DG GABAergic neurons which inhibit GCs. We took advantage of specific methods and a gain- and loss-of function strategy with Designer Receptors Exclusively Activated by Designer Drugs (DREADDs) to study MCs in diverse behaviors. Using this approach, manipulations of MCs could bidirectionally regulate behavior. The results suggest that inhibiting MCs can reduce anxiety-like behavior and improve cognitive performance. However, not all cognitive or anxiety-related behaviors were influenced, suggesting specific roles of MCs in some but not all types of cognition and anxiety. Notably, several behaviors showed sex-specific effects, with females often showing more pronounced effects than the males. We also used the immediate early gene c-Fos to address whether DREADDs bidirectionally regulated MC or GC activity. We confirmed excitatory DREADDs increased MC c-Fos. However, there was no change in GC c-Fos, consistent with MC activation leading to GABAergic inhibition of GCs. In contrast, inhibitory DREADDs led to a large increase in GC c-Fos, consistent with a reduction in MC excitation of GABAergic neurons, and reduced inhibition of GCs. Taken together, these results suggest that MCs regulate anxiety and cognition in specific ways. We also raise the possibility that cognitive performance may be improved by reducing anxiety.

**SIGNIFICANCE STATEMENT:** The dentate gyrus (DG) has many important cognitive roles as well as being associated with affective behavior. This study addressed how a glutamatergic DG cell type called mossy cells (MCs) contributes to diverse behaviors, which is timely because it is known that MCs regulate the activity of the primary DG cell type, granule cells (GCs), but how MC activity influences behavior is unclear. We show, surprisingly, that activating MCs can lead to adverse behavioral outcomes, and inhibiting MCs have an opposite effect. Importantly, the results appeared to be task-dependent and showed that testing both sexes was important. Additional experiments indicated what MC and GC circuitry was involved. Taken together, the results suggest how MCs influence behaviors that involve the DG.

## 1. INTRODUCTION

The dentate gyrus (DG) is critical to hippocampal function and is also implicated in psychiatric disorders (Scharfman, 2007b). Dentate granule cells (GCs) are the primary excitatory cell type in the DG and receive input from cortical regions such as the entorhinal cortex. GCs represent the first component of the trisynaptic circuit (GCs→CA3→CA1) and are therefore essential for propagating information throughout the hippocampus. Within the DG, GCs are regulated by GABAergic inhibitory neurons and glutamatergic hilar mossy cells (MCs). MCs are in a unique position to regulate GC activity because they project directly to GC dendrites (MC→GC), but also indirectly inhibit GCs through their innervation of local GABAergic neurons (MC→GABAergic neuron→GC). The complex circuitry of MCs in the DG has led to extensive debate about their net effects on GCs (Ratzliff et al., 2002; Sloviter et al., 2003; Jinde et al., 2013; Scharfman, 2016, 2017).

Several studies have suggested that MCs are important for spatial functions of the DG (Soltesz et al., 1993; Danielson et al., 2017; GoodSmith et al., 2017; Senzai and Buzsaki, 2017; GoodSmith et al., 2019). A limited number of studies have also shown that MCs influence other DG functions, such as contextual discrimination and object learning (Jinde et al., 2012; Bui et al., 2018; Azevedo et al., 2019). MCs have also been implicated in recognizing novelty in the environment such as the presence of new objects (Bernstein et al., 2019). Moreover, MCs are sensitive to restraint stress (Moretto et al., 2017), which is interesting because of studies linking the DG to affective behaviors, including anxiety (McEwen et al., 2016; Anacker and Hen, 2017). However, there remains a limited understanding about the role of MCs in anxiety-like behaviors. Part of this uncertainty is due to conflicting reports about MCs in anxiety-like behaviors from previous studies (Jinde et al., 2012; Bui et al., 2018; Oh et al., 2019), possibly attributable to the different methods in targeting and manipulating MCs. In addition, the majority of MC studies to date have focused on male subjects (Jinde et al., 2012; Duffy et al., 2013; Moretto et al., 2017; Senzai and Buzsaki, 2017; Oh et al., 2019) or did not provide a clear view of sex differences (Danielson et al., 2017; GoodSmith et al., 2017; Bui et al., 2018). The focus on male subjects is problematic because there are known sex differences in GC structure, activity and synaptic plasticity (Hajszan et al., 2007; Zitman and Richter-Levin, 2013; Harte-Hargrove et al., 2015; Yagi and Galea, 2019) and some data showing sex differences in MCs (Guidi et al., 2006).

To clarify the role of MCs in cognitive and anxiety-like behaviors, we used a gain- and loss-of function approach using Designer Receptors Exclusively Activated by Designer Drugs (DREADDs) in female and male mice. Remarkably, inhibition of MCs benefited cognitive and anxiety-related behaviors in several tasks, especially those associated with objects in an environment, which could be interpreted as contextual cues. In contrast, excitation of MCs was generally associated with adverse behavioral effects. We also used c-Fos as a tool to understand how DREADDs modified the activity of MCs and GCs. Excitatory DREADDs (eDREADDs) increased MC but not GC activity, supporting the view that MCs primarily inhibit GCs by activating intermediary GABAergic neurons. Conversely, inhibitory DREADDs (iDREADDs) approximately doubled the number of active GCs, consistent with reduced inhibition of GCs through the MC→GABAergic neuron→GC pathway. Notably, several behavioral tasks showed female- or male-specific DREADD effects, indicating that both sexes are necessary to avoid an underestimation of effects on the DG. Taken together, our results suggest that lowering MC activity can benefit both cognitive and anxiety-related behavior.

Therefore, MCs are an important cell type in cognitive and anxiety-like behaviors.

## 2. METHODS

### 2.1 Terminology

It is acknowledged that the use of the term anxiety for a mouse is difficult to distinguish from fear or behavioral stress (Bailey and Crawley, 2009; LeDoux and Pine, 2016; Fanselow and Pennington, 2017). In many parts of the text we use ‘anxiety-like’ to reflect the importance of being cautious about the use of the term anxiety.

### 2.2 Experimental design and controls

All experimental procedures were completed in accordance with the National Institutes of Health (NIH) guidelines and approved by the Institutional Animal Care and Use Committee at the Nathan Kline Institute. The present study used transgenic Drd2-Cre^+/-^ mice to selectively target and manipulate the activity of MCs *in vivo* using excitatory and inhibitory DREADDs. Importantly, electrophysiological studies from our lab (Botterill et al., 2019) and others (Yeh et al., 2018; Oh et al., 2019) have confirmed the excitatory and inhibitory effects of DREADDs in MCs. Control mice consisted of Drd2-Cre^-/-^ and Cre^+/-^ mice injected with a viral control fluorophore (mCherry). Mice recovered for 3 weeks after surgery to allow for viral expression and then underwent a series of behavioral tests to evaluate the role of MCs in cognitive and anxiety-like behaviors. Each behavioral test was spaced at least one week apart, except for three anxiety tests that were done on the same day. These tests were the open field test (OFT), light-dark box (LDB), and elevated plus maze (EPM). These tests were done on the same day because our prior experience suggested they did not influence each other. The order of the behaviors were: week 1, OFT, LDB, EPM; week 2, novel object location (NOL); week 3, novel object recognition (NOR); week 4, novelty suppressed feeding (NSF); week 5, contextual fear conditioning (CFC), week 6, home cage novel object exploration (HCNOE).

Mice were acclimated to handling by experimenters to minimize stress associated with repeated handling and injections. DREADDs were activated with clozapine-N-oxide (CNO, 5 mg/kg, i.p., #BML-NS105-0005, Enzo Life Sciences) one hour prior to behavioral testing unless noted otherwise below. The dose of 5mg/kg CNO was selected because it is reported to robustly activate DREADDs with minimal off-target behavioral effects reported at higher doses (MacLaren et al., 2016; Manvich et al., 2018). Control mice were also injected with CNO to further control for potential off-target effects. After behavioral testing was completed, mice were euthanized, and brain tissue was prepared for immunohistochemical analyses to evaluate viral expression and immediate early gene (IEG) activity, as described below. Unless noted otherwise, behavioral scores pertaining to time were measured in seconds (sec) and distance in meters. Statistical comparisons were made using tests and criteria defined below.

### 2.3 Animals and genotyping

Male and female Drd2-Cre transgenic mice (8-18 weeks old) maintained on a C57BL/6N background were used for all experiments. Breeding was done in house as previously described (Botterill et al., 2019). Mice were weaned at postnatal day 25-30 and housed with same-sex siblings in standard laboratory cages (2-4 per cage) with corn cob bedding. Mice were maintained on a 12 hour light-dark cycle with standard rodent chow (Purina 5001, W.F. Fisher) and water available *ad libitum*. Genotyping was performed by the Genotyping Core Laboratory at New York University Langone Medical Center.

### 2.4 Viral targeting of mossy cells

To target MCs and their axons that span the septotemporal extent of the DG, virus was injected bilaterally into the rostral and caudal hippocampus as previously described (Botterill et al., 2019). Drd2-Cre^+/-^ mice were injected with eDREADDs (AAV2-hSyn-DIO-hM3D(Gq)-mCherry; ≥5×10^12^ vg/mL, #44361, Addgene) or iDREADDs (AAV5-hSyn-DIO-hM4D(Gi)-mCherry; ≥8×10^12^ vg/mL, #44362, Addgene; Figure 1A). Controls were injected with a mCherry construct (AAV5-EF1a-DIO-mCherry; ≥3×10^12^ vg/mL, University of North Carolina Vector Core).

**Figure 1.**
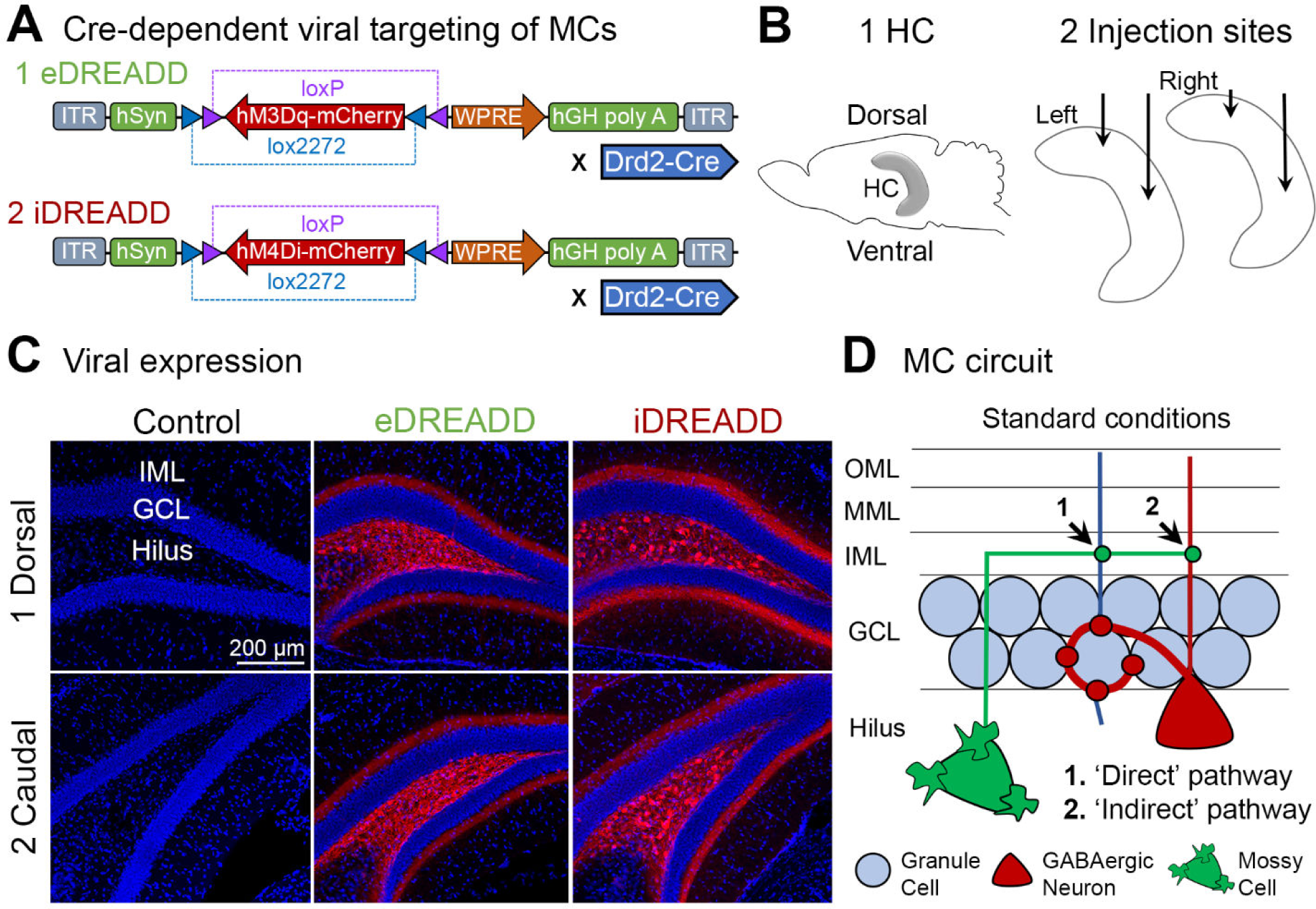
Experimental design. **(A)** Viral constructs used for **(A1)** gain-of-function (excitatory DREADD; eDREADD) and **(A2)** loss-of-function (inhibitory DREADD; iDREADD) experiments. **(B1)** Schematic of the hippocampus. **(B2)** 160nL of virus was injected into the rostral and caudal hippocampus, bilaterally. **(C)** Representative viral expression in **(C1)** dorsal and **(C2)** caudal coronal sections of control, eDREADD, and iDREADD mice. Inner molecular layer (IML), granule cell layer (GCL). Scale bar: 200µm. **(D)** Simplified MC circuit diagram. *(1)* MCs excite GCs through a monosynaptic ‘direct’ pathway. *(2)* MCs also inhibit GCs through an ‘indirect’ MC→GABAergic neuron→GC inhibitory pathway. The indirect inhibitory pathway is thought to dominate the direct excitatory pathway under normal conditions.

### 2.5 Stereotaxic surgery and viral injections

Stereotaxic surgery was performed as described previously (Botterill et al., 2019). Briefly, mice were anesthetized with isoflurane (5 % induction, 1-2 % maintenance; Aerrane, Henry Schein) and secured in a rodent stereotaxic apparatus (Model #502063, World Precision Instruments). Buprenex (Buprenorphine, 0.1 mg/kg, s.c.) was delivered prior to surgical procedures to reduce discomfort. Body temperature was maintained at 37 °C via a homeothermic blanket system (Harvard Apparatus). The scalp of each mouse was shaved and swabbed with betadine (Purdue Products) and lubricating gel was applied to the eyes to prevent dehydration (Patterson Veterinary).

A surgical drill (Model C300, Grobert) was used to make craniotomies bilaterally over the rostral (−2 mm anterior-posterior, ±1.2 mm medial-lateral) and caudal hippocampus (−3.2 mm anterior-posterior, ±2.3 mm medial-lateral), relative to bregma. A 500 nL Neuros Syringe (#65457-02, Hamilton Company) attached to the stereotaxic apparatus was positioned over each craniotomy and lowered 2.0 mm (rostral) or 2.6 mm (caudal) below the skull surface for viral delivery (Figure 1B). Each of the 4 injection sites was injected with 160 nL of virus at a rate of 80 nL per minute. The needle remained in place for at least 5 minutes after the injection to allow for diffusion of the virus and then the needle was slowly removed from the brain. After all viral injections were complete, the scalp of each mouse was cleaned with sterile saline and sutured using tissue adhesive (Vetbond, 3M). Mice were transferred to a clean cage at the end of the surgery and placed on a heating blanket (37 °C) until fully ambulatory.

### 2.6 Behavioral tests

All behavioral tests were conducted in dedicated procedure rooms. All testing arenas were made in house and the dimensions are provided below. Mice remained in their home cage after the CNO injection until behavioral testing. At the end of each behavioral test, mice were returned to their home cage and left undisturbed until the next test. For all experiments, the testing arenas and equipment were cleaned thoroughly with 70 % ethanol (EtOH) between subjects.

All behavioral tests were recorded with a Logitech C920 1080P webcam connected to a PC (Logitech® Webcam software, v. 2.51). Experimenters blinded to the experimental conditions manually reviewed and scored the behavioral tests offline. ANY-maze tracking software (v. 6.2; Stoelting Co., USA) was used to score the OFT, LDB, and EPM tests. Notably, we manually scored a subset of these videos and found a Pearson correlation coefficient of r=0.99 with the ANY-maze scores.

#### 2.6.1 CFC

CFC was conducted as previously described with minor modifications (Stone et al., 2011). Briefly, mice were placed inside a Plexiglas fear conditioning chamber (18 cm x 18 cm x 20 cm) placed inside of a larger arena (34 cm x 45 cm x 34 cm). The floor of the fear conditioning chamber contained 28 stainless steel rods (0.2 cm diameter, spaced 0.5 cm apart). Mice were placed inside the Plexiglas chamber and allowed to acclimate for 2 minutes. After the baseline period, 3 foot shocks (0.5 mA for 2 sec) were delivered once per minute. The mice remained in the fear conditioning chamber for an additional 2 minutes after the final foot shock (4 minutes total) and were then returned to their home cage. Contextual fear memory was assessed 24 hours later by placing mice into the same chamber where training occurred for 10 minutes. Freezing behavior was operationally defined as the termination of all motor movements except those necessary for respiration (Fanselow, 1980; Botterill et al., 2015a; Botterill et al., 2015b; Guskjolen et al., 2018). Data are reported as percent freezing, calculated by dividing the time spent freezing (sec) each minute for RMANOVA analyses or by dividing the total time freezing (sec) by test duration (i.e., 240 sec training and 600 sec testing) for average freezing scores.

#### 2.6.2 NOR and NOL

To evaluate the role of MCs on spatial and object memory, mice underwent the NOR and NOL tests as previously described with minor modifications (Leger et al., 2013; Vogel-Ciernia and Wood, 2014; Brymer et al., 2020). Briefly, both tasks involve presenting two identical objects during a training session and evaluating object exploration during a subsequent test session. The difference between the two tasks is that one of the two previously presented objects is an entirely new object replaces one of the training objects during the NOR test, whereas in the NOL test, one of the two identical objects is moved to a new location (Vogel-Ciernia and Wood, 2014). Both tasks are based on the premise that rodents have an innate preference for novelty (e.g., a novel object or moved object). Importantly, both the NOR and NOL tests are thought to involve the DG (Kinnavane et al., 2015; Kesner, 2018).

##### 2.6.2.1 Acclimation & training

For both tasks, mice underwent 3 acclimation sessions (5 minutes each day) prior to training (day 4). Each acclimation session consisted of a brief handling session followed by placing the mouse in a rectangular testing arena (24 cm x 45 cm x 20 cm) that was located inside of a large arena (40 cm x 62 cm x 46 cm) with visual cues on each wall. Mice were injected with CNO 30 minutes before the training session and then placed in the same rectangular cage described above and allowed to explore two identical novel objects (“A” & “B”) spaced 5 cm apart for 5 minutes. Each training session introduced an identical pair of Legos (3 cm x 4.5 cm x 5 cm) or bronze pineapples (3.5 cm diameter, 5.5 cm tall) that were secured to the base of the testing arena. Mice were returned to their home cage after completing the training session.

##### 2.6.2.2 Testing

One hour after the training session, mice were returned to the rectangular testing arena and allowed to explore for 5 minutes. For the NOR test, one familiar object from the training session was replaced with a novel object (object “B”) spaced 5 cm from object “A”. In the NOR test, novel object “B” was a 20 mL scintillating vial (2.5 cm diameter x 6 cm tall) filled with an opaque gel. For the NOL test, object “A” remained in the same location as training, but object “B” was moved approximately 20 cm to the other side of the testing arena.

##### 2.6.2.3 Analysis

For both the training and testing procedures, the amount of time mice spent exploring each object was measured. The preference for the novel or moved object “B” in the NOR and NOL test was determined by calculating an object discrimination index; [DI = (T_*B*_ – T_*A*_) / (T_*B*_ + T_*A*_)] *100, where T_*B*_ represents time spent exploring object “B” and T_*A*_ represents time spent exploring object “A”. Mice were considered to explore an object when their head was facing the object and the nose was approximately within 1 cm of the object. Mice that failed to explore objects during training (i.e., less than 1 sec) were removed from the analysis, similar to criteria reported elsewhere (Bui et al., 2018).

#### 2.6.3 HCNOE

To evaluate the role of MCs on object exploration in a familiar environment, we used a modified version of the HCNOE test recently described by our laboratory (Bernstein et al., 2019). At least 3 days prior to testing, mice were transferred into a clean cage and allowed to acclimate to the behavioral testing room. On the test day, mice were injected with CNO 90 minutes prior to testing. Two identical novel objects (Legos: 3 cm x 4.5 cm x 5 cm) were placed in the home cage, spaced approximately 15 cm apart and 5 cm from the cage walls. Mice were allowed to explore the two objects for a total of 10 minutes. We used the same criteria for object exploration as described for NOL and NOR tests. The percent of time spent exploring objects was calculated by the amount of time (sec) exploring objects each minute for RMANOVA analyses or by summing the total time exploring objects and dividing it by 240 sec (first 4 minutes). Mice were sacrificed 90 minutes after completing the test to evaluate immunoreactivity of the IEG c-Fos (see section 2.7 below).

#### 2.6.4 NSF

To evaluate whether MCs contribute to feeding behaviors in a novel environment, mice underwent the NSF test as previously described with minor modifications (Dulawa and Hen, 2005; Demireva et al., 2018). Briefly, mice were food deprived for 24 hours and water deprived for 2 hours prior to the start of the test. At the start of each session, the mouse was placed in the corner of a brightly illuminated novel arena (51 cm x 51 cm x 17 cm) and allowed to explore for 10 minutes. A rodent chow pellet was placed in the middle of the open field arena. The latency to feed was measured, defined as the interval between placing the mouse in the chamber and the time to begin eating the chow pellet. Mice that did not feed during the test received a maximum score of 600 sec.

#### 2.6.5 LDB

Mice were tested in the LDB which is designed to probe the innate aversion of rodents to brightly illuminated areas (Klemenhagen et al., 2006; Takao and Miyakawa, 2006). Mice were placed in a chamber containing a brightly illuminated light compartment and a dimly lit dark compartment of equal size (20 cm x 20 cm x 22 cm). The light and dark compartments were connected through an open partition (7 cm wide x 7 cm high) that allowed the mice to freely move throughout the two chambers. At the start of each test, mice were placed in the center of the arena facing the dark compartment. Mice were removed from the testing arena after 5 minutes. Anxiety-like and locomotor behaviors were evaluated by measuring the time spent in the light compartment, the latency to enter the light compartment, and the distance traveled in the light compartment.

#### 2.6.6 OFT

We also evaluated exploratory and anxiety-like behaviors in the OFT (Seibenhener and Wooten, 2015; Teixeira et al., 2018). Mice were placed in the periphery of a brightly illuminated open field (42 cm x 42 cm x 30 cm) and allowed to explore the arena for 10 minutes and then returned to their home cage. Anxiety-like behavior was assessed by measuring the time spent in the center of the open field (24 cm x 24 cm). Locomotor behavior was assessed by measuring the total distance traveled during the task.

#### 2.6.7 EPM

The EPM was used to test exploratory and anxiety-like behavior (Komada et al., 2008). The EPM apparatus consisted of two open and closed arms of identical dimensions (5 cm x 22 cm). The closed arms had 15 cm high walls whereas the open arms had 3 mm high ledges to prevent mice from falling off the apparatus. Arms of the same type were arranged at opposite sides to each other and were raised 55 cm above the floor. At the start of each test, the mouse was placed in the central square (6 cm x 6 cm) of the EPM apparatus facing one of the closed arms. Mice were allowed to explore the apparatus for 5 minutes. The measures of interest were the percent of time spent in the open arms of the apparatus which was determined by calculating time spent in the open arms (sec) divided by test duration (300 sec), the number of open arm entries, and the total distance traveled during the task.

### 2.7 Anatomy

#### 2.7.1 Perfusion-fixation and sectioning

Mice were initially anesthetized with isoflurane, followed by urethane (2.5 g/kg; i.p.). Once under deep anesthesia, the abdominal cavity was opened and the subject was transcardially perfused with ∼10 mL of room temperature saline, followed by ∼20 mL of cold 4 % paraformaldehyde in 0.1 M phosphate buffer (PB; pH =7.4). The brains were extracted and stored overnight at 4 °C in 4 % paraformaldehyde in 0.1 M PB. The brains were then hemisected and sectioned in the coronal (right hemisphere) or horizontal (left hemisphere) plane at 50 µm (Vibratome 3000, Ted Pella). Sections were collected using a 1 in 12 series (600 µm apart). For subsequent analyses, we used at least 3 sections for each region of interest (e.g., rostral vs caudal and dorsal vs ventral measurements). To evaluate the dorsal hippocampus, sections were cut in the coronal plane because it maintains the lamination of the DG well. For the ventral hippocampus, where coronal sections make the different parts of the DG hard to interpret, sections were cut in the horizontal plane. Sections were stored in 24-well tissue culture plates containing cryoprotectant (30 % sucrose, 30 % ethylene glycol in 0.1 M PB) at -20 °C until use.

#### 2.7.2 Viral expression

The expression of hM4D(Gi) or hM3D(Gq) in Drd2-Cre^+/-^ mice was visualized by the mCherry tag (Figure 1C). Viral expression in Drd2-Cre^+/-^ mice was characterized by large hilar mCherry^+^ cells proximal to the injection site and a dense band of mCherry^+^ labeling in the inner molecular layer (IML) throughout the septotemporal axis of the DG, consistent with the location of MCs and their major axon projection (Figure 1C-D; Scharfman, 2016). The pattern of viral expression has been validated in previous work by our laboratory (Botterill et al., 2019; Bernstein et al., 2020) and confirmed by others (Danielson et al., 2017; Bui et al., 2018; Yeh et al., 2018; Azevedo et al., 2019; Oh et al., 2019).

Briefly, sections were rinsed in 0.1 M Tris Buffer (TB, 3 x 5 minutes), followed by 0.1 M TB containing 0.25 % Triton X-100 (Tris A), and 0.1M TB containing 0.25 % Triton X-100 and 1 % bovine serum albumin (Tris B). The sections were blocked with 5 % normal goat serum in Tris B for 30 minutes and incubated overnight at 4 °C with a rabbit polyclonal primary antibody against mCherry (1:3000, #167453, Abcam) diluted in blocking solution. On the following day, the sections were incubated with goat anti-rabbit Alexa Fluor 568 secondary antibody (1:1000, #A11036, Invitrogen) in Tris B. The sections were counterstained with Hoechst 33342 (1:20000), mounted onto microscope slides, and coverslipped with Citifluor (Electron Microscopy Sciences) mounting medium. Images were acquired with a LSM 880 laser scanning confocal microscope (Zeiss) using a 10 x objective and frame size of 2048 x 2048 pixels. Any mouse that was injected with virus encoding DREADDs that lacked viral expression (due to mistargeted injections or incorrect genotype) was removed from the study.

#### 2.7.3 C-Fos immunoreactivity

Mice were euthanized 90 minutes after completing HCNOE (180 minutes after CNO) to evaluate the effect of DREADDs on c-Fos immunoreactivity. We examined c-Fos after HCNOE because we have previously reported that c-Fos is effective in staining active MCs and GCs following HCNOE (Bernstein et al., 2019). Sections spaced approximately 600 µm apart were rinsed in 0.1M TB (3 x 5 minutes) followed by 1 % H_2_0_2_ in 0.1 M TB for 5 minutes to block endogenous peroxidase activity. Sections were then rinsed in Tris A and Tris B (10 minutes each) and then incubated for 30 minutes in 5 % (v/v) normal goat serum diluted in Tris B (blocking solution). The sections were then incubated overnight at 4 °C in rabbit polyclonal anti-c-Fos primary antibody (1:2000, #226003, Synaptic Systems) diluted in blocking solution. This antibody is widely used and highly specific for c-Fos protein (Zhou et al., 2019; Kim and Cho, 2020). On the following day, sections were rinsed in 0.1 M TB (3 x 5 minutes) and incubated in biotinylated goat anti-rabbit secondary antibody (1:500, Vector) diluted in Tris B for 2 hours. The sections were then rinsed in 0.1 M TB (2 x 5 minutes) and incubated in avidin-biotin complex (1:500, #PK-6100 VECTASTAIN Elite, Vector) for 1 hour. Sections were visualized by incubating them in a solution containing 0.5 mg/mL 3,3’-diaminobenzidine tetrahydrochloride (Sigma), 40 µg/mL ammonium chloride (Sigma), 25 mg/mL (D+)-glucose (Sigma), and 3 µg/mL glucose oxidase (Sigma) in 0.1 M TB. The reaction was halted by rinsing sections in 0.1 M TB (3 x 5 minutes). Sections were mounted on gelatin-coated slides and dried overnight at room temperature. On the following day, the sections were dehydrated using a graded EtOH series (70 %, 95 %, 100 %), cleared in Xylene, and coverslipped with Permount (Electron Microscopy Sciences). Photomicrographs were captured using a 10 x objective on an Olympus BX61 microscope equipped with a CCD camera (Retiga 2000R, QImaging).

#### 2.7.4 C-Fos quantification

We analyzed c-Fos immunoreactivity across the septotemporal axis of the DG using criteria previously reported by our laboratory (Duffy et al., 2013; Moretto et al., 2017; Bernstein et al., 2019). Immunoreactive cells were manually counted at 16 x at similar locations across the septotemporal axis between subjects as previously described (Botterill et al., 2014; Moretto et al., 2017). The total number of c-Fos immunoreactive cells in the hilus and GCL were divided by the number of sections to determine the average number of cells per section.

### 2.8 Data analysis and statistics

All results are presented as mean ± standard error of the mean (SEM). For all analyses, statistical significance was achieved if the *p* value was <0.05 (denoted on all graphs by an asterisk). Statistical comparisons were conducted in Prism 8.4 (GraphPad).

For parametric data with multiple comparisons, two-way ANOVAs were performed. When a statistically significant main effect was observed (e.g., treatment or sex), post-hoc tests (Tukey’s or Sidak) were used with corrections for multiple comparisons. When the main effect of treatment (e.g., control, eDREADD, iDREADD) was significant, main effects within the female and male cohorts were analyzed using the above mentioned post-hoc tests. When the interaction of factors was not significant, it was not reported in the Results. For all data sets, the ROUT method (Prism) was used to detect and remove outliers using nonlinear regression. When Bartlett’s tests showed that the variance of groups was not equal, data were transformed using a log10 function. Notably, the statistical results of transformed data were similar to the raw data. Statistical values for the transformed data are reported in the Results. Graphs show raw data.

Sample sizes were determined with power analysis (G*Power software). We determined that for a two-tailed analysis with significance set at α = 0.05 and power > 80%, approximately 8-10 subjects per treatment were required. For all analyses, at least 10 subjects per treatment were used when sex was pooled. We acknowledge some of the data sets have less than 10 subjects per treatment when evaluating male and female differences and this could impact statistical power. However, several analyses within the male and female cohorts detected treatment differences with as few as 5-6 subjects, suggesting that the study was adequately powered.

### 2.9 Additional technical considerations

This study targeted most MCs. However, the observed effects may have been more robust if all MCs expressed DREADDs. On the other hand, activating all MCs may lead to different effects than activating only those that are dorsal or ventral. In addition, there could be different effects in a different background strain or species. Regarding females, we did not examine effects of the estrous cycle. One of the reasons is that females that are stressed usually have irregular estrous cycles, and our study involved stressors (e.g., CNO injections). In addition, it is important to bear in mind that there are considerable sex differences in the response to stress in rodents (Luine et al., 2007; Bale and Epperson, 2015). Other the other hand, other studies in mice and rats have found some of the effects we observed, such as sex differences in exploration and cognition (Galea et al., 2017; Yagi and Galea, 2019). Regardless, the results suggest we think is very important, that restricting studies to males may underestimate the role of the DG in some experiments and overestimate it in others.

## 3. RESULTS

### 3.1 Behavioral tests

#### 3.1.1 CFC

Given the importance of the DG in contextual learning and memory (Phillips and LeDoux, 1992), we were interested to determine whether MCs contribute to CFC. Our primary measurement was conditioned freezing, as defined in the Methods. Notably, CNO was administered prior to training, but not testing.

##### 3.1.1.1 CFC Training

We first measured freezing behavior during the training session (Figure 2A). Baseline (B) freezing (time points B1, B2 in Figure 2B) and post-shock (PS) freezing (PS1 through PS4 in Figure 2B) were evaluated on a minute by minute basis for the training session. Note that only 3 shocks were delivered during training, so PS4 represents the final minute of the training session. A two-way RMANOVA found a main effect of time (*F*(5,195)=41.96, *p*<0.001) and a time by treatment interaction (*F*(10,195)=2.185, *p*=0.020). Tukey’s post-hoc test revealed that there was no effect of treatment on baseline freezing (all *p* values >0.612; Figure 2B). Similarly, there was no effect of treatment on freezing behavior on PS1 or PS2 (all *p* values >0.051; Figure 2B). However, Tukey’s post-hoc test revealed that control mice (28.69 ± 4.95 %) engaged in significantly greater freezing behavior than eDREADD mice (14.45 ± 2.71 %; *p*=0.017) in the minute after the third shock (PS3; Figure 2B). By the final minute of the training session (PS4), both control (34.96 ± 5.19 %) and iDREADD mice (38.43 ± 5.86 %) engaged in approximately twice as much freezing behavior as eDREADD mice (19.99 ± 3.27 %; all *p* values <0.011; Figure 2B).

**Figure 2.**
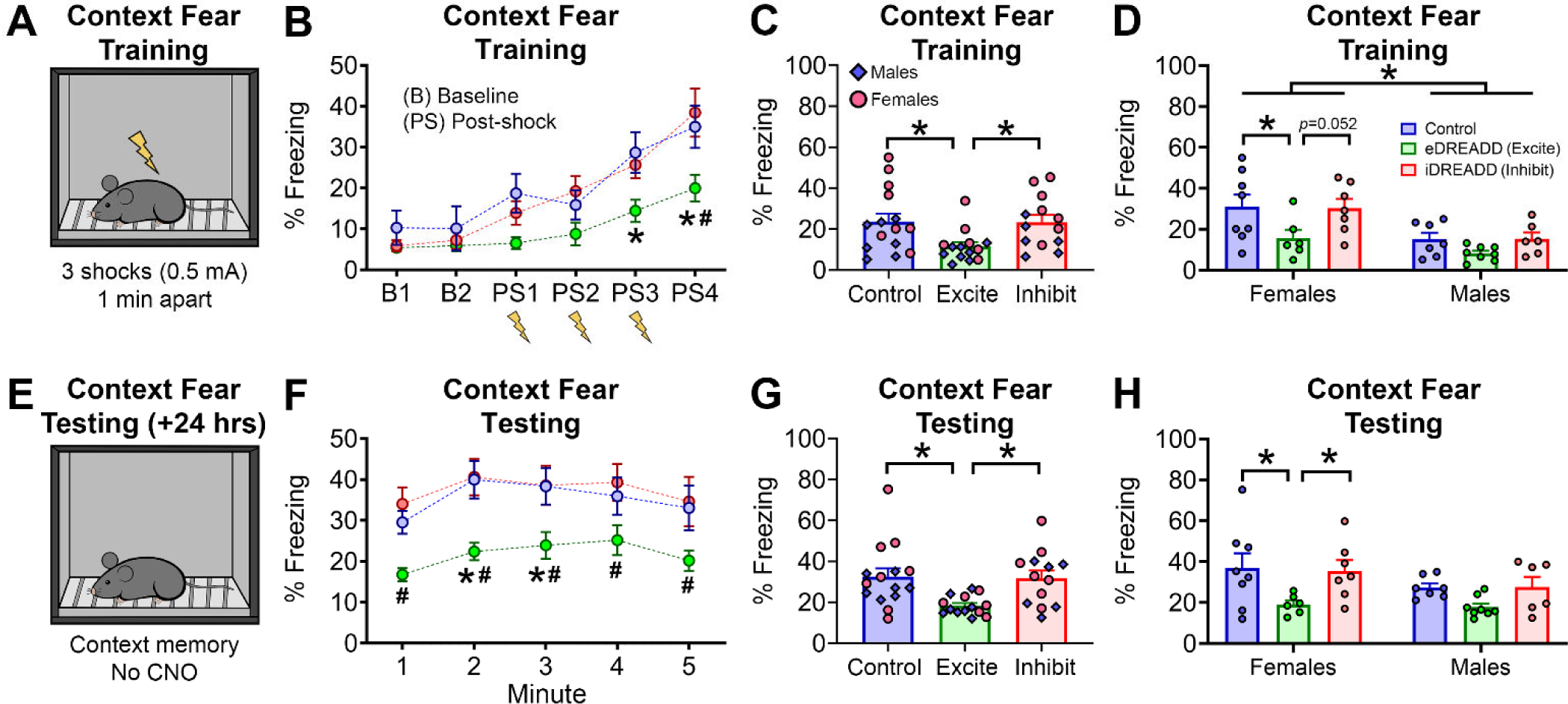
CFC in control, eDREADD and iDREADD mice. **(A)** Mice were placed in a fear conditioning chamber and 3 footshocks (0.5mA) were delivered 1 minute apart. **(B)** Minute by minute analysis of the training session found no effect of treatment on baseline freezing (B1 & B2) or freezing during the first 2 post-shock minutes (PS1 & PS2). The eDREADD mice froze significantly less than controls in the third post-shock minute (PS3; *p*=0.017) and less than control and iDREADD mice in the fourth minute (PS4; all *p* values <0.011). **(C)** When data were averaged across all post-shock minutes, eDREADD mice froze significantly less than control and iDREADD mice (all p values <0.037). **(D)** Female eDREADD mice froze significantly less than female control mice (*p*=0.036) and female iDREADD mice had a similar pattern (*p*=0.052). There was a sex difference in training, with female mice freezing significantly more than male mice (*p*<0.001). Also, there was no significant effect of treatment in the male cohort. **(E)** Mice were returned to the same fear conditioning apparatus 24 hours later to assess fear memory. **(F)** Minute by minute analysis of the first 5 minutes of the context test showed that eDREADD mice froze less than iDREADD (all *p* values <0.047) and control mice (all *p* values <0.033). **(G)** When freezing behavior was averaged across the entire test duration, eDREADD mice showed significantly less freezing than control and iDREADD mice (all *p* values <0.011). **(H)** There was a significant effect of treatment in the female cohort, whereby eDREADD mice froze significantly less than control and iDREADD mice (all *p* values <0.043). There was no effect of treatment in the male cohort. Data are represented as mean ± SEM. * denotes *p*<0.05. In panels **B & F, \*** denotes significantly different from control (*p*<0.05), while **#** denotes iDREADD significantly different from eDREADD (*p*<0.05).

In Figure 2C, the average post-shock freezing across the 4 minutes of the training session is shown. A two-way ANOVA revealed an overall effect of treatment (*F*(2,36)=4.711, *p*=0.015). Tukey’s post-hoc test found that the total time freezing during the training session was significantly greater in iDREADD (23.34 ± 3.51 %) and control mice (23.59 ± 3.98 %) compared to eDREADD mice (11.46 ± 2.08 %; all *p* values <0.037; Figure 2C). Control and iDREADD mice did not differ (*p*=0.997).

In the two-way ANOVA, sex was also a significant main factor (*F*(1,36)=14.43, *p*<0.001). Figure 2D separates female and male data to compare the data more easily. Notably, there was a greater percent of freezing in females (26.38 ± 3.22 %) than males (12.56 ± 1.57 %; Figure 2D). Also, Tukey’s post-hoc test showed that female control mice (30.98 ± 5.97 %) froze significantly more than female eDREADD mice (15.70 ± 4.12 %; p=0.036). Female iDREADD mice showed a similar pattern of freezing behavior as control mice (30.27 ± 4.60 %), but they did not statistically differ from eDREADD mice (*p*=0.052). Interestingly, male control, eDREADD and iDREADD mice did not differ in freezing behavior during training (all *p* values >0.443), suggesting the female mice were primarily driving the treatment differences observed during training (Figures 2B-D). The higher freezing scores in female mice are consistent with a recent study that reported females show greater fear generalization and freezing (Keiser et al., 2017).

##### 3.1.1.2 CFC Testing

Mice were tested for contextual fear memory 24 hours after training by placement in the same chamber without a shock (Figure 2E). Minute-by-minute comparisons are shown in Figure 2F and pooled data from all 10 minutes of the test are shown in Figure 2G. Minute-by-minute data, analyzed with a RMANOVA, showed a significant effect of treatment (*F*(2,39)=6.033, *p*=0.005) and a significant effect of time (*F*(4,156)=5.314, *p*<0.001). Tukey’s post-hoc test found that eDREADD mice froze significantly less than iDREADD mice for each of the first 5 minutes of the memory test (all *p* values <0.047). Furthermore, eDREADD mice froze significantly less than control mice in the second and third minute of the memory test (all p values <0.033). Note that greater freezing during the memory test is considered a reflection of better recall of the noxious stimulus delivered during the training session.

Using pooled data (Figure 2G), a two-way ANOVA found a significant effect of treatment (*F*(2,36)=6.731, *p*=0.003), with Tukey’s post-hoc test finding greater freezing in iDREADD mice (31.84 ± 3.70 %) and control mice (32.53 ± 4.01 %) compared to eDREADD mice (18.39 ± 1.27 %; all *p* values <0.011; Figure 2G). Freezing behavior in iDREADD mice and controls did not differ significantly (*p*=0.989). There was no difference between male and female cohorts during testing, indicated by no main effect of sex (*F*(1,36)=2.013, *p*=0.164). Female and male data are shown separately in Figure 2H. Tukey’s post-hoc test indicated that freezing behavior during the memory test was significantly greater in female control (37.04 ± 7.12 %) and iDREADD mice (35.49 ± 5.30 %) relative to eDREADD mice (19.15 ± 1.94 %; all *p* values <0.043; Figure 2H). There was no difference between treatments in male mice during the memory test (all *p* values > 0.094).

Overall, the data suggest that eDREADD treatment worsened performance both during the learning phase and memory phase of the task.

#### 3.1.2 NOR

A recent study suggests that information about objects acquired in the lateral EC (LEC) from sensory input may influence MCs because the LEC projects to MCs (Azevedo et al., 2019). Therefore, we evaluated object recognition memory using the NOR task (Figure 3A).

**Figure 3.**
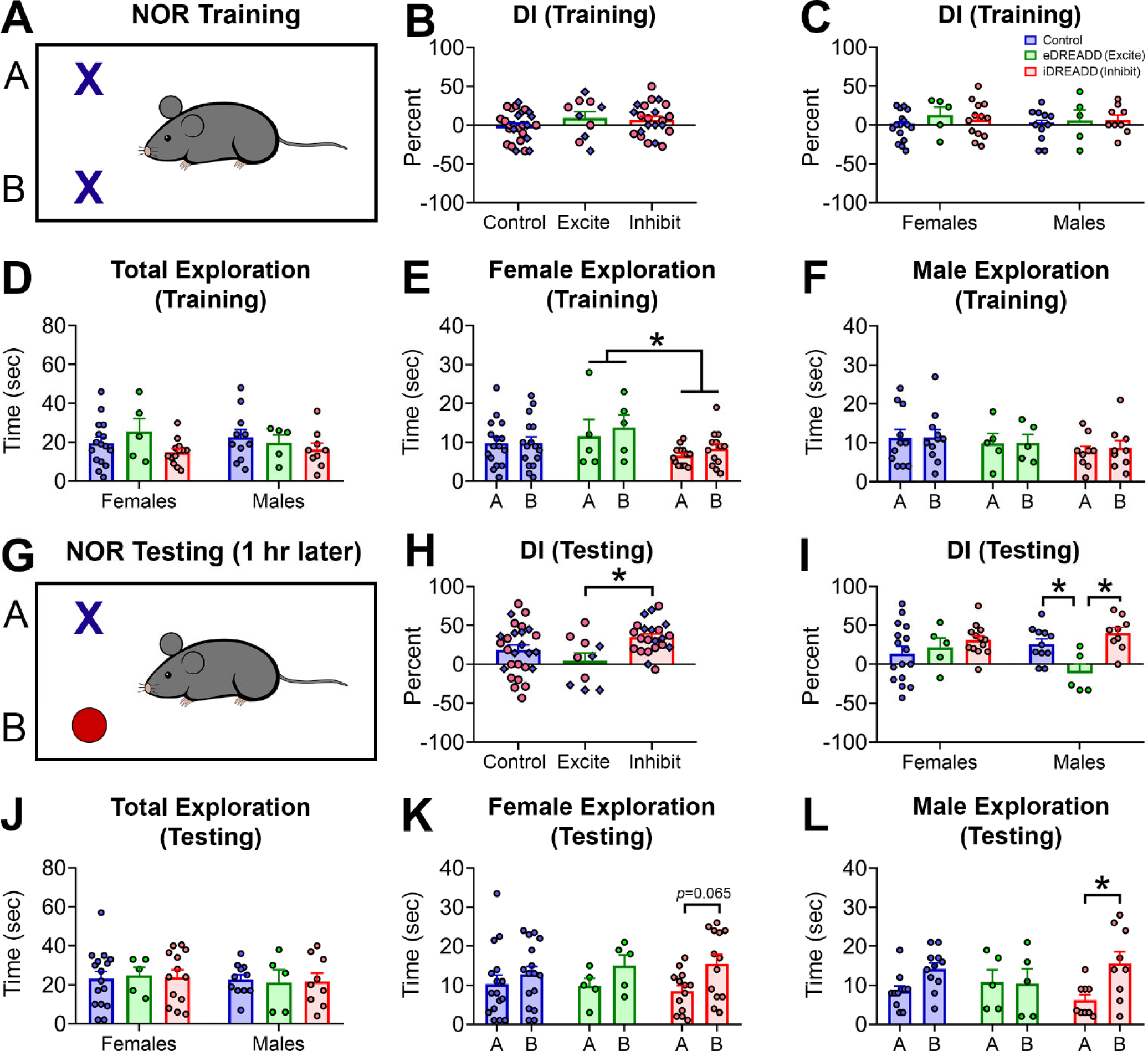
NOR in control, eDREADD and iDREADD mice. **(A)** In NOR training, mice explored two identical novel objects for 5 minutes. **(B)** There was no effect of treatment on the training discrimination index (DI) when both sexes were pooled. **(C)** There was no effect of treatment on training DI in male and female cohorts. **(D)** There was no effect of treatment on the total time spent exploring objects (“A” + “B”) during NOR training in female and male cohorts. **(E)** Female iDREADD mice spent significantly less time exploring objects than female eDREADD mice during NOR training (*p*=0.044). **(F)** Male mice did not differ by treatment on time spent exploring object “A” versus “B” during training. **(G)** Mice were tested for object recognition memory one hour after training by replacing object “B” with a novel object. **(H)** iDREADD mice had a significantly greater testing DI than eDREADD mice (*p*=0.013). **(I)** Testing DI did not differ by treatment in female mice. However, male control and iDREADD mice had a significantly greater testing DI than eDREADD mice (all *p* values <0.034). **(J)** Female and male mice did not differ by treatment in total object exploration during testing. **(K)** There was no effect of treatment in female mice on the time spent exploring object “A” versus “B” during testing (all *p* values >0.065). **(L)** Male iDREADD mice spent significantly more time exploring object “B” than “A” during testing (*p*=0.008).

##### 3.1.2.1 NOR Training

###### A) Discrimination index

First, we calculated the DI during training by comparing the amount of time spent exploring object “A” versus object “B” (see Methods). A two-way ANOVA found that the training DI did not differ by treatment (*F*(2,53)=1.159, *p*=0.321) or sex (*F*(1,53)=0.099, *p*=0.753; Figure 3B-C).

###### B) Total exploration time

Next, we evaluated the total time spent exploring objects during training (i.e., “A” + “B”). A two-way ANOVA found no effect of treatment (*F*(2,53)=2.018, *p*=0.143), or sex (*F*(1,53)=0.017, *p*=0.894; Figure 3D) on total object exploration.

###### C) Object exploration time

Next, we evaluated object exploration time, meaning the time in seconds that objects “A” and object “B” were explored. These data reduce the data in the training DI and total exploration time to the raw data for each object. In female mice, a two-way ANOVA with treatment and object as factors showed a significant effect of treatment (*F*(2,62)=3.188, *p*=0.048) but not object (*F*(1,62)=0.744, *p*=0.391) on exploration. Tukey’s post-hoc test showed that exploration by female eDREADD mice was greater than female iDREADD mice (*p*=0.044; Figure 3E). In male mice, there were no effects of treatment on time spent exploring object “A” versus “B” (*F*(1,44)=0.065, *p*=0.799; Figure 3F). Although these data suggest female eDREADD mice explored slightly more than female iDREADD mice during training, the results also show that there was no effect of treatment on object preference during training. This is an important distinction because any preference for one object during training makes the results of testing hard to interpret (Vogel-Ciernia and Wood, 2014).

##### 3.1.2.2 NOR Testing

###### A Discrimination index

Object recognition memory was evaluated 1 hour after training by replacing object “B” of the training session with a novel object “B” (Figure 3G). A two-way ANOVA found a significant effect of treatment (*F*(2,53)=4.636, *p*=0.013), but no effect of sex (*F*(1,53)=0.280, *p*=0.598). Tukey’s post-hoc test showed that iDREADD mice had a significantly greater testing DI (34.77 ± 4.38 %) than eDREADD mice (4.85 ± 9.80 %; *p*=0.013; Figure 3H). The iDREADD mice were not significantly different from control mice (18.66 ± 6.11 %; all *p* values >0.127), which were between eDREADD and iDREADD mice (Figure 3H).

To further investigate the treatment effect, we analyzed effects within female and male cohorts. Notably, Tukey’s post-hoc test found that the testing DI in male eDREADD mice (−11.94 ± 11.97 %) was significantly lower than control (25.88 ± 6.58 %) and iDREADD mice (40.06 ± 7.22 %; all *p* values <0.034; Figure 3I). This result is consistent with worse performance in eDREADD mice. The results from female mice showed greater variability than males on testing DI and this is likely to have contributed to the lack of a treatment difference between control, eDREADD and iDREADD mice (all *p* values >0.212).

###### B Total exploration time

Next, we evaluated the total time spent exploring objects “A” and “B” during testing. A two-way ANOVA found no effect of treatment (*F*(2,53)<0.001, *p*=0.999) or sex (*F*(1,53)=0.336, *p*=0.564; Figure 3J).

###### C Object exploration time

In males, there was no effect of treatment (*F*(2,44)=0.052, *p*=0.949), but a significant difference in the time spent exploring object “A” versus “B” (*F*(1,44)=6.77, *p*=0.012) during testing. Sidak’s multiple comparisons test found that male iDREADD mice spent significantly more time exploring object “B” (15.56 ± 3.02 sec) than object “A” (6.16 ± 1.39 sec; *p*=0.008; Figure 3L). There were no differences between object “A” versus “B” in male control or eDREADD mice (all *p* values >0.112).

For female mice, a two-way ANOVA found no effect of treatment (*F*(2,62)=0.052, *p*=0.949), but a significant difference in time spent exploring object “A” versus “B” (*F*(1,62)=5.454, *p*=0.022). Thus, females appeared to have a slight preference for the novel object, independent of treatment. This preference was small, however, and in support of this interpretation, Sidak’s multiple comparisons test showed none of the paired comparisons were significantly different (all *p* values >0.065; Figure 3K).

These data suggest that inhibiting MCs led to improved object recognition memory. Both males and females showed the effect, but statistical comparisons were significant only for males. Taken together, the data suggest that inhibiting MCs can benefit cognitive performance in NOR.

#### 3.1.3 NOL

To evaluate object location memory, mice underwent the NOL task (Figure 4A). First, we evaluated the training DI and a two-way ANOVA found no effect of treatment (*F*(2,53)=0.276, *p*=0.759) or sex (*F*(1,53)=0.288, *p*=0.593; Figure 4B-C). However, there was a statistically significant interaction (*F*(2,53)=5.337, *p*=0.007), whereby control, eDREADD, and iDREADD mice showed a pattern of opposing DI scores in their respective female and male cohorts (Figure 4C).

**Figure 4.**
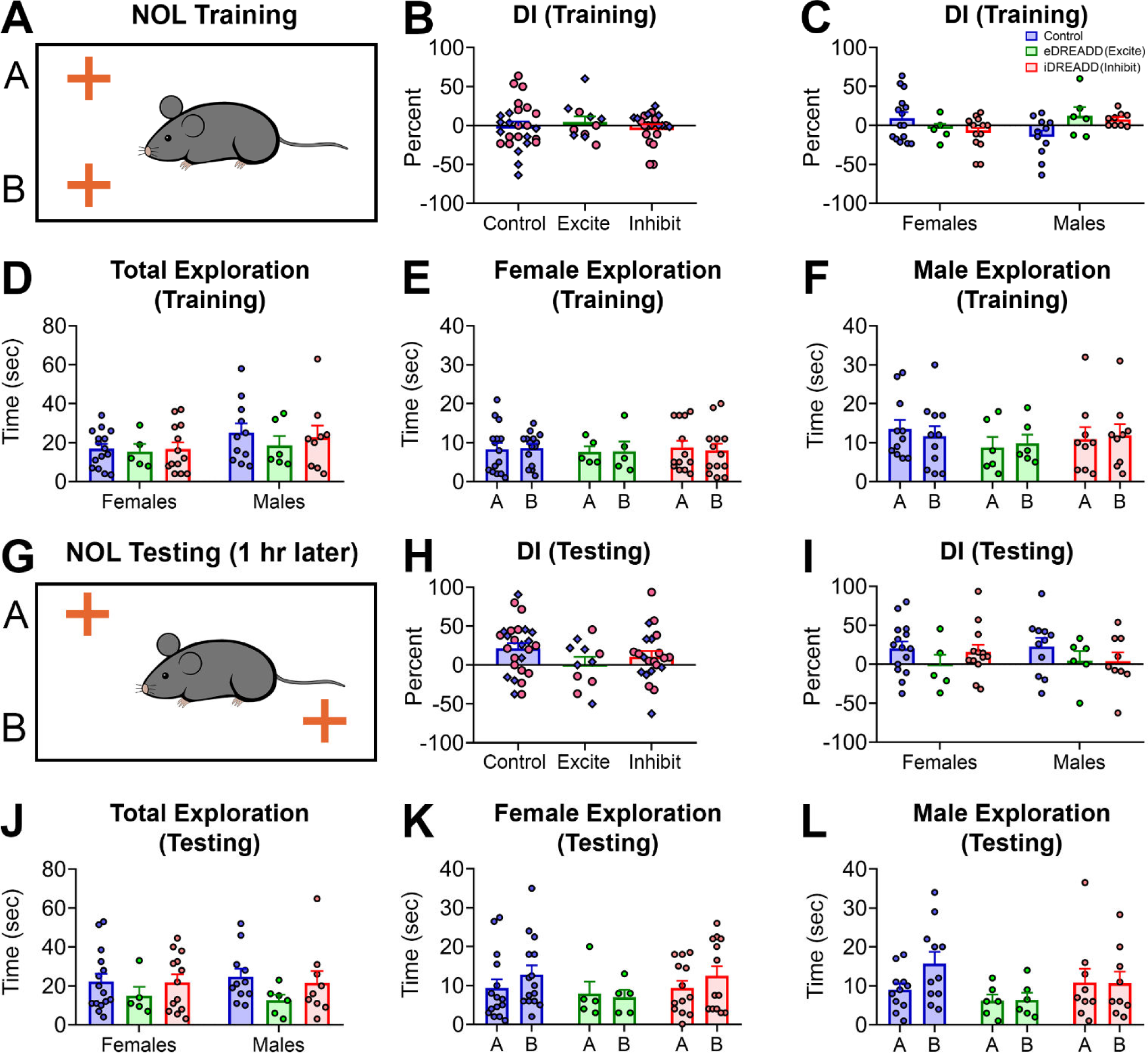
NOL in control, eDREADD and iDREADD mice. **(A)** In NOL training, mice explored two identical novel objects for 5 minutes. **(B)** The overall NOL training discrimination index (DI) did not differ by treatment. **(C)** There was no effect of treatment on NOL training DI in the female and male cohorts. **(D)** Female and male mice did not differ in total object exploration time (“A” + “B”) during training. **(E-F)** Female and male mice did not differ by treatment in the time spent exploring object “A” versus “B” during training. **(G)** Mice were tested for object location memory one hour later by moving object “B” to the other side of the testing arena. **(H)** There was no significant effect of treatment on the testing DI. **(I)** The testing DI did not differ by treatment in male and female cohorts. **(J)** Female and male mice did not differ in their total object exploration time during testing. **(K-L)** There was no effect of treatment in female and male mice on spent time spent exploring object “A” versus “B” during testing.

Next, we measured the total amount of time exploring both objects (i.e., “A” + “B”) and a two-way ANOVA found no effect of treatment (*F*(2,53)=0.355, *p*=0.702) or sex (*F*(1,53)=2.438, *p*=0.124; Figure 4D). Furthermore, there were no effects of treatment on time spent exploring object “A” versus “B” in female (*F*(1,60)=0.002, *p*=0.959) or male mice (*F*(1,46)=0.001, *p*=0.969; Figure 4E-F).

Object location memory was tested 1 hour later during the test phase by moving object “B” to the other side of the testing arena (Figure 4G). A two-way ANOVA found that treatment had no significant effect on the testing DI (*F*(2,53)=1.622, *p*=0.207) and sex did not either (*F*(1,53)=0.006, *p*=0.935; Figure 4H-I).

A two-way ANOVA also revealed that there was no effect of treatment on the total time spent exploring both objects during testing (*F*(2,53)=1.743, *p*=0.184), and there was no effect of sex either (*F*(1,53)<0.001, *p*=0.992; Figure 4J). Furthermore, there was no effect of treatment in the amount of time spent exploring object “A” versus “B” in female (*F*(1,60)=0.701, *p*=0.405) or male mice (*F*(1,46)=0.976, *p*=0.328; Figure 4K-L).

In summary, there appeared to be little effect of treatment in the NOL task. However, there are several potential reasons for the lack of an effect in NOL (see Discussion).

#### 3.1.4 HCNOE

Next, we used the HCNOE task (Figure 5A), which we have found activates MCs in a robust manner, but not many other cells in the DG or hippocampus (Duffy et al., 2013; Bernstein et al., 2019). Interestingly, this task involves the home cage to reduce behavioral stress, so it is highly relevant to the present study.

**Figure 5.**
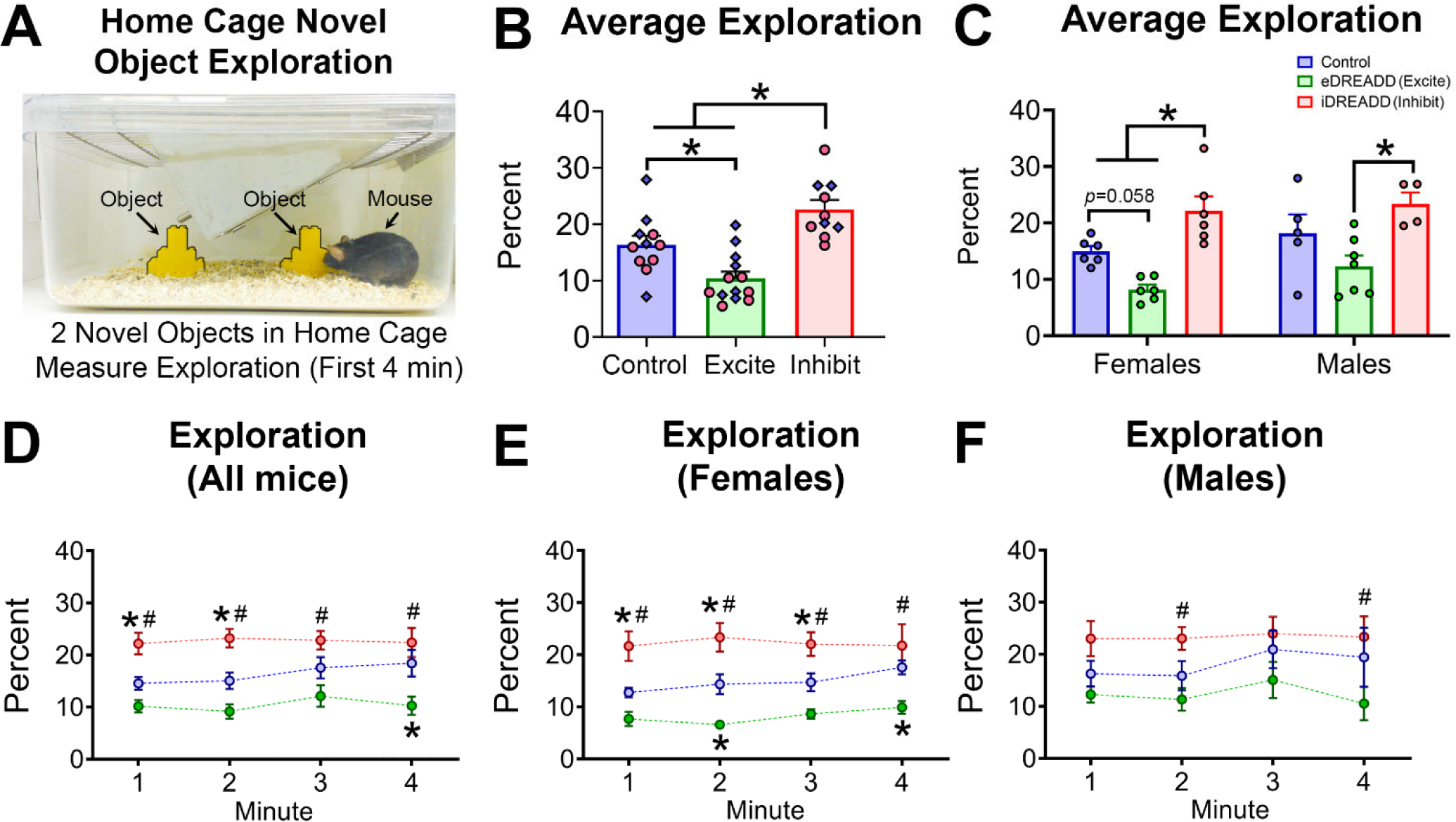
HCNOE in control, eDREADD and iDREADD mice. **(A)** Two identical novel objects (yellow Legos, outlined in black; see arrows) were placed in the home cage. Object exploration was measured over the first 4 minutes. **(B)** There was an overall effect of treatment on object exploration, with iDREADD mice spending a greater percent of time exploring objects than control and eDREADD mice (all *p* values <0.019). Furthermore, eDREADD mice spent less time exploring objects compared to control mice (*p*=0.010). **(C)** There was a significant effect of treatment in the female cohort, with iDREADD mice spending a greater percent of time exploring than control and eDREADD mice (all *p* values <0.039). Also, male iDREADD mice spent a greater time exploring objects than male eDREADD mice (*p*=0.003). **(D)** Minute by minute analysis found that iDREADD mice spent a greater percent of time exploring than eDREADD mice for each of the 4 minutes (all *p* values <0.001) and greater exploration than control mice for the first 2 minutes (all *p* values <0.017). Control mice also showed a greater percent of exploration than eDREADD mice during the fourth minute (*p*=0.005). **(E-F)** Minute by minute exploration in female and male mice. Overall, similar effects were observed as in the pooled analysis shown in **D**. Thus, iDREADD mice generally showed greater exploration than eDREADD mice and controls were often between the two treatment groups. In panels **D-F, \*** denotes significantly different from control (*p*<0.05), while **#** denotes iDREADD significantly different from eDREADD (*p*<0.05).

##### 3.1.4.1 Average exploration

First, we focused on the percent of time exploring objects during the first 4 minutes of HCNOE. A two-way ANOVA found a significant effect of treatment (*F*(2,28)=18.32, *p*<0.001), but not sex (*F*(1,28)=2.755, *p*=0.108). Tukey’s post-hoc test reporting that iDREADD mice (22.66 ± 1.64 %) spent significantly more time exploring objects than control mice (16.40 ± 1.59 %) and eDREADD mice (10.41 ± 1.21 %; all *p* values <0.019; Figure 5B). Conversely, eDREADD mice spent significantly less time exploring objects compared to control mice (*p*=0.010), consistent with worse performance described in other tasks above.

The data for each sex are plotted separately in Figure 5C and show the similarities between the female and male cohorts on HCNOE exploration. Tukey’s post-hoc test showed that some pairwise comparisons were significant, similar to the pooled data in Figure 5C. For example, female iDREADD mice spent significantly more time exploring (22.20 ± 2.51 %) than female control mice (14.95 ± 0.92 %) and eDREADD mice (8.19 ± 0.84 %; all *p* values <0.039; Figure 5C). Female control and female eDREADD mice did not differ from each other although the *p* value approached criterion (*p*=0.058). For males, iDREADD mice spent a greater percent of time exploring objects (23.35 ± 2.03 %) than eDREADD mice (12.31 ± 1.91 %; *p*=0.003). The male control mice (18.14 ± 3.35 %) scored between iDREADD and eDREADD mice and did not differ significantly (all *p* values >0.120).

##### 3.1.4.2 Exploration minute by minute

Next, we analyzed object exploration over each of the first 4 minutes of the HCNOE task (Figure 5D). A two-way RMANOVA revealed an overall effect of treatment (*F*(2,31)=17.57, *p*<0.001) but not time (*F*(3,93)=1.341, *p*=0.265). Tukey’s post-hoc tests revealed that iDREADD mice showed a greater percent of time exploring than eDREADD mice for each of the 4 minutes (all *p* values <0.001; Figure 5D). The iDREADD mice also showed a greater percent of exploration than control mice for the first 2 minutes of the analysis (all *p* values <0.017). Finally, the control mice showed a greater percent of exploration than eDREADD mice on the fourth minute of the task (*p*=0.005).

When each sex was examined separately, a two-way RMANOVA in female mice found a significant effect of treatment (*F*(2,15)=18.34, *p*<0.001) but not time (*F*(3,45)=1.353, *p*=0.269). Tukey’s post-hoc test revealed that for the first 3 minutes of the test, female iDREADD mice showed a greater percent of exploration than control and eDREADD mice (all *p* values <0.039; Figure 5E). In the fourth minute, female iDREADD mice were significantly different than eDREADD mice (*p*<0.001). Moreover, female eDREADD mice spent a lesser percent of time exploring objects than control mice during minutes 2 and 4 (all *p* values <0.029). These data show a robust effect of treatment in females.

In male mice, a two-way RMANOVA revealed a significant effect of treatment (*F*(2,13)=4.884, *p*=0.026) but not time (*F*(3,39)=1.353, *p*=0.269). Tukey’s post-hoc test found treatment differences in the second and fourth minute of the test, with iDREADD mice spending a greater percent of time exploring objects than eDREADD mice at both times (all *p* values <0.042; Figure 5F). These data suggest a similar effect of treatment in males as females, but effects in males were not as robust because all minutes of the session did not show treatment differences.

In summary, iDREADDs significantly improved performance in the HCNOE task, and conversely, eDREADDs worsened performance, consistent with several of the prior tasks.

#### 3.1.5 NSF

NSF is commonly used to evaluate aversion to eating in a brightly illuminated, novel environment (Figure 6A). In light of a recent study suggesting that MCs may regulate feeding behavior (Azevedo et al., 2019), it was timely to use this test to gain further insight into effects of MC on behavior.

**Figure 6.**
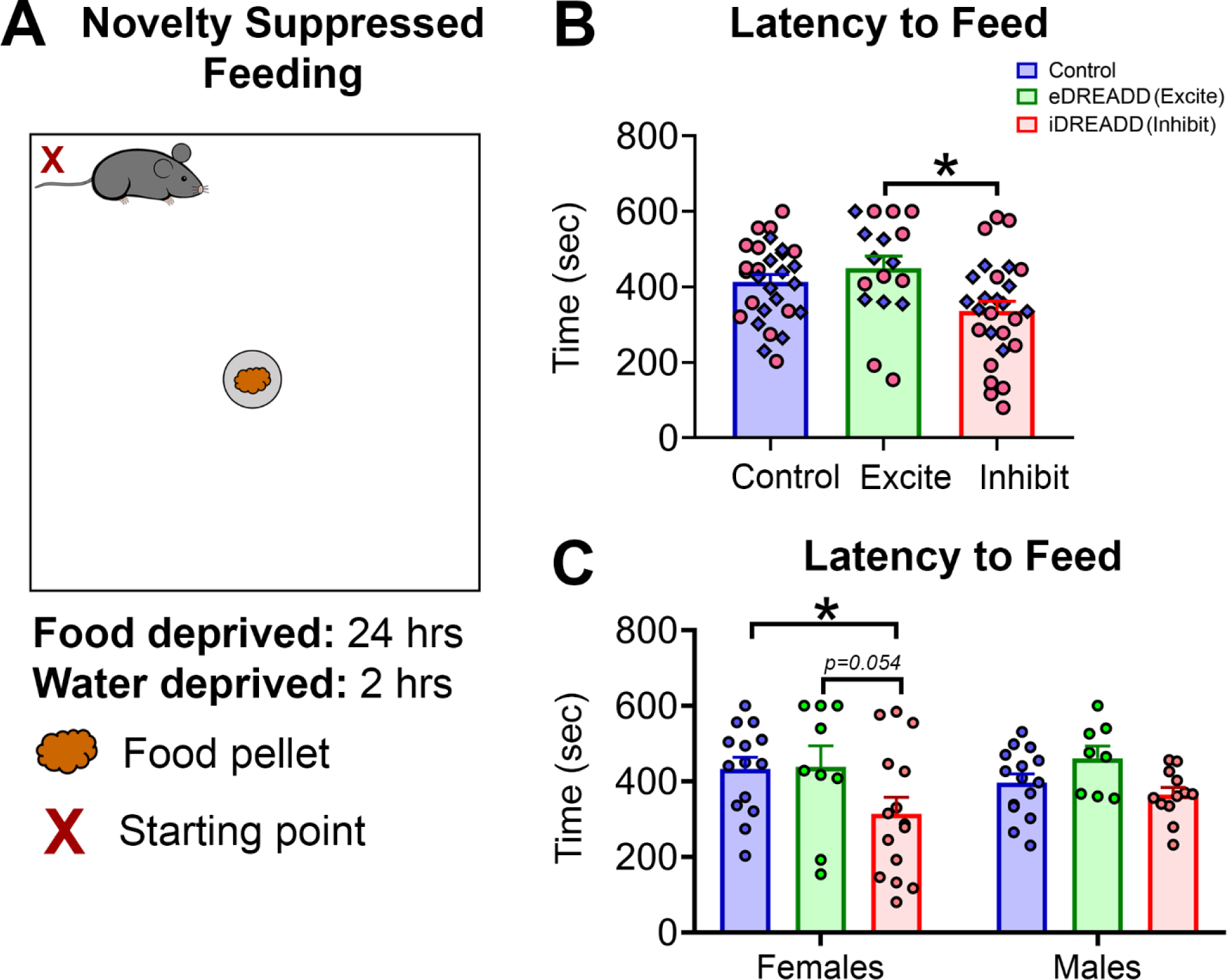
NSF in control, eDREADD and iDREADD mice. **(A)** Mice were food deprived for 24 hours and water deprived for 2 hours before undergoing the NSF test. Mice were placed in the corner of a brightly illuminated novel arena (“X”) and the latency to eat a food pellet in the arena was measured. **(B)** There was a significant effect of treatment, with iDREADD mice eating approximately 30% sooner than the eDREADD mice (*p*=0.015). There were no other treatment differences in latency to feed. **(C)** Female iDREADD mice had a significantly shorter latency to feed compared to control mice (*p*=0.033). No other significant treatment differences were found between female mice (all *p* values >0.054). The latency to feed did not differ between treatments in male mice.

A two-way ANOVA revealed a significant main effect of treatment (*F*(2,67)=4.652, *p*=0.012) but no effect of sex (*F*(1,67)=0.187, *p*=0.666) on the latency to feed. Tukey’s post-hoc test showed that iDREADD mice (336.4 ± 25.92 sec) had a shorter latency to feed than eDREADD mice (448.7 ± 32.90 sec; *p*=0.015; Figure 6B). No other comparisons showed a significant treatment difference in the latency to feed (all *p* values >0.069).

The data from females and males are shown in Figure 6C. Tukey’s post-hoc test found that female iDREADD mice engaged in feeding behavior significantly sooner (313.9 ± 44.0 sec) than control female mice (432.1 ± 31.39 sec; *p*=0.033; Figure 6C). A similar pattern was seen when comparing the female iDREADD and eDREADD mice (Figure 6C) but this effect did not reach criterion (*p*=0.054). There was no significant effect of treatment in the male mice (all *p* values >0.213).

In summary, inhibiting MCs had an effect consistent with reduced anxiety-like behavior. These data are also consistent with the recent observation that iDREADD treatment in Drd2-Cre mice facilitates feeding behaviors (Azevedo et al., 2019).

#### 3.1.6 LDB

Next, we evaluated anxiety-like behavior associated with the natural aversion of mice to a brightly illuminated area in the LDB (see Methods).

A two-way ANOVA revealed a significant effect of treatment on the percent of time mice spent in the light compartment (*F*(2,60)=3.525, *p*=0.035). Tukey’s post-hoc test found the iDREADD mice spent approximately 25% more time in the light compartment (77.35 ± 6.12 sec) than control mice (60.38 ± 3.70 sec; *p*=0.036; Figure 7A), consistent with an anxiolytic effect. The main effect of sex was not significant (*F*(1,60)=1.027, *p*=0.315), suggesting that the female and male cohorts showed similar behaviors in the LDB. To further investigate the effect of treatment, we evaluated simple main effects within female and male cohorts. The male iDREADD mice spent a greater percent of time in the light compartment (80.84 ± 8.32 sec) compared to male control mice (57.46 ± 5.93 sec *p*=0.037; Figure 7B-C). Several of the female iDREADD mice also appeared to spend more time in the light compartment, similar to the iDREADD males, but there were no statistical differences in the female cohort (all *p* values >0.185; Figure 7B). Locomotor activity was quantified as the total distance traveled within the lighted compartment. A two-way ANOVA showed that there were no significant effects of treatment (*F*(2,60)=0.946, *p*=0.394; Figure 7D) or sex (*F*(1,60)=0.103, *p*=0.749; Figure 7E). There also was no effect of treatment (*F*(2,60)=0.294, *p*=0.746), or sex on the latency to enter the light compartment (*F*(1,60)=1.498, *p*=0.225; data not shown).

**Figure 7.**
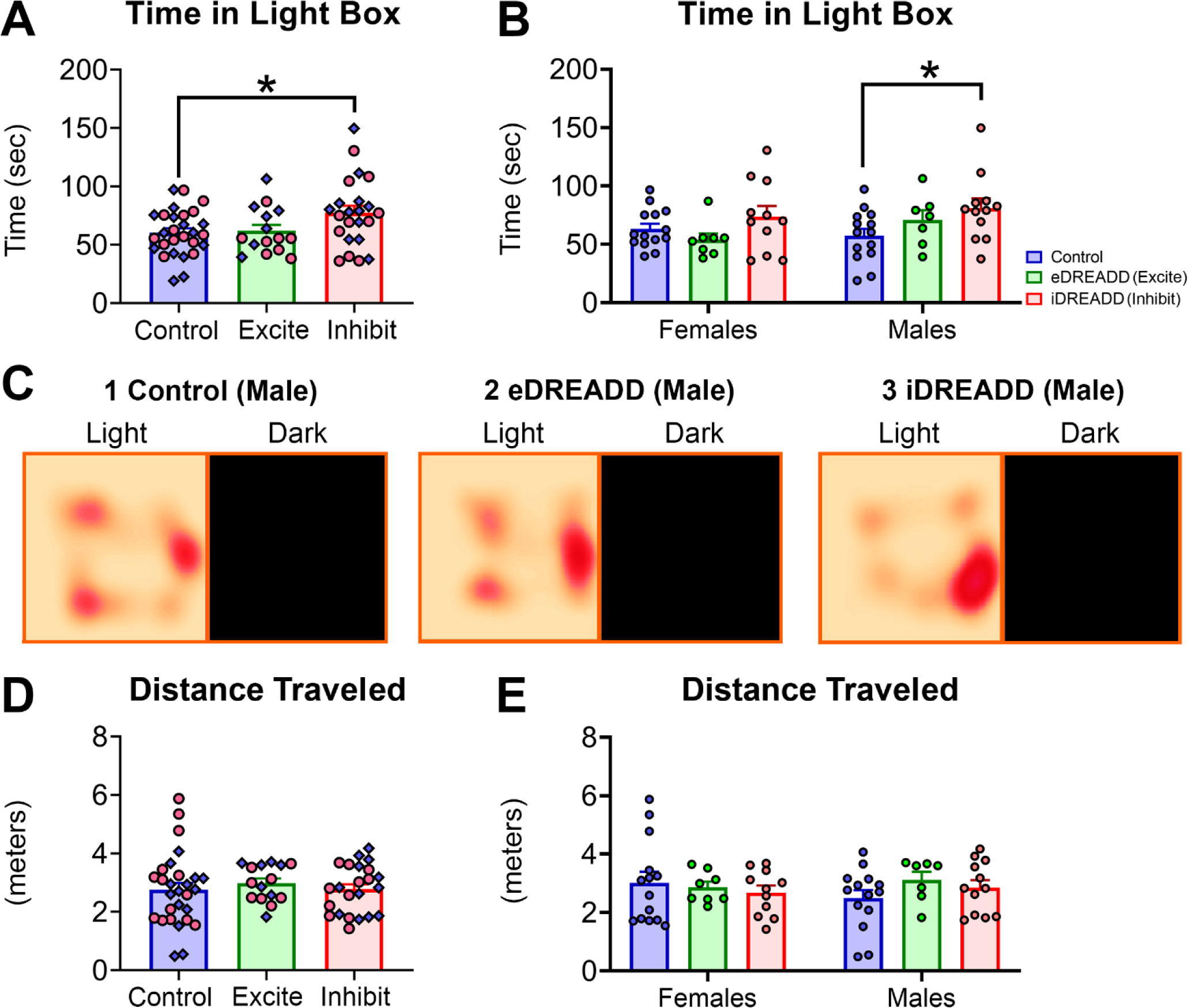
LDB in control, eDREADD and iDREADD mice. **(A)** iDREADD mice spent approximately 25% more time in the light compartment of the LDB compared to control mice (*p*=0.036). **(B)** There was no effect of treatment in female mice on the amount of time spent in the light compartment of the LDB. However, male iDREADD mice spent more time in the light compartment of the LDB compared to male control mice (*p*=0.037). **(C)** Representative heat maps of male **(C1)** control, **(C2)** eDREADD, and **(C3)** iDREADD mice in the light compartment of the LDB. **(D-E)** There was no effect of treatment on the distance traveled in the light compartment of the LDB when subjects were pooled or separated by sex.

In summary, LDB results suggest an anxiolytic effect of inhibiting MCs with males showing a more robust effect than females.

#### 3.1.7 OFT

In the OFT, the time spent in the center of the open field was analyzed using a two-way ANOVA with treatment and sex as factors. There was no effect of treatment (*F*(2,67)=2.616, *p*=0.080; Figure 8A), but there was a significant effect of sex (*F*(1,67)=6.768, *p*=0.011) attributable to female mice spending approximately 25% less time (68.25 ± 5.89 sec) in the center of the open field than the male mice (89.06 ± 4.79 sec; Figure 8B). These data suggest females showed more anxiety-like behavior than males, an idea that has been discussed extensively before in humans (Donner and Lowry, 2013; Altemus et al., 2014), but depends on several factors in rodents (Palanza, 2001; Simpson and Kelly, 2012).

**Figure 8.**
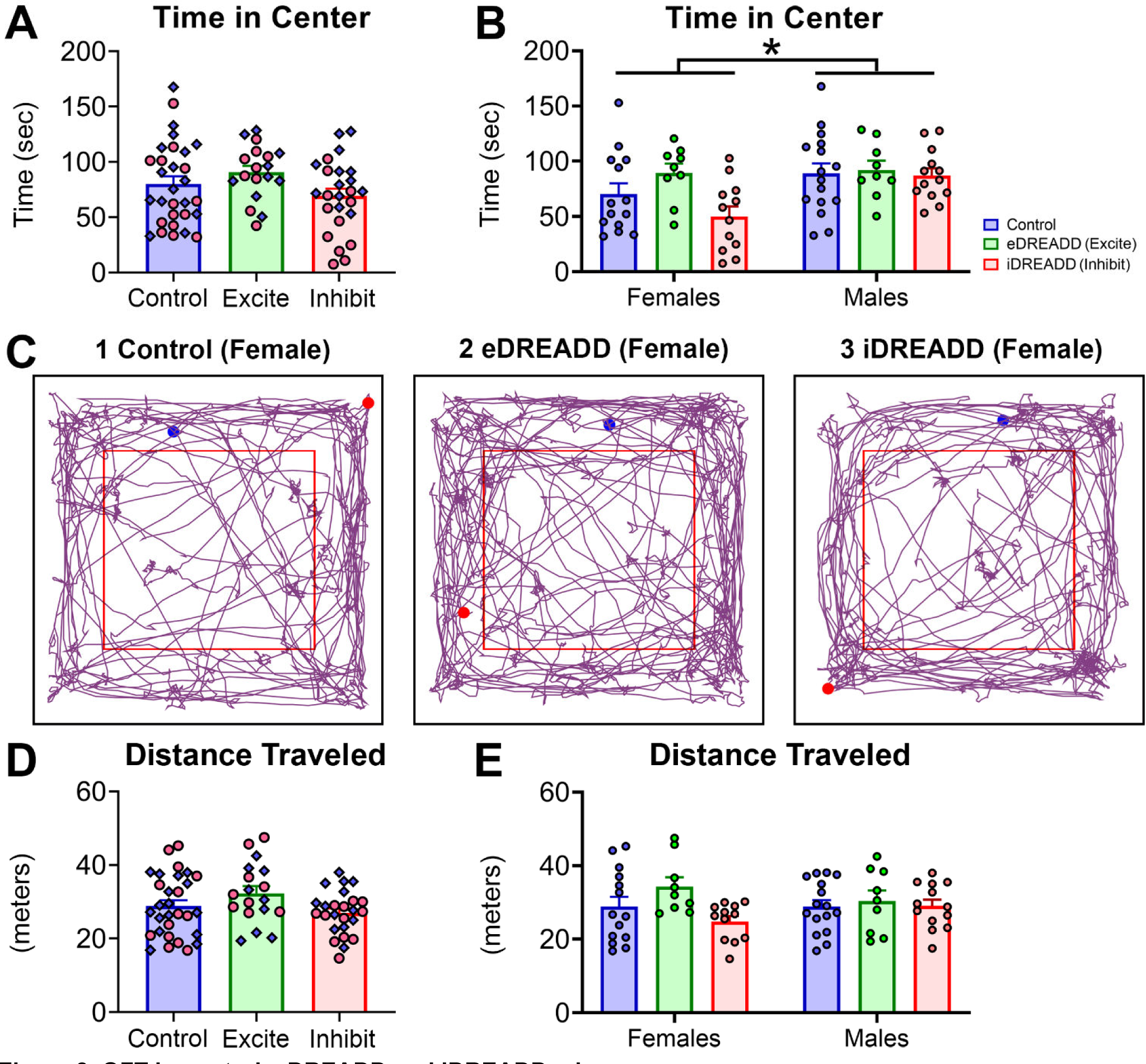
OFT in control, eDREADD and iDREADD mice. **(A)** DREADD treatment had no significant effect on the amount of time spent in the center of the OFT. **(B)** Female mice spent significantly less time in the center of the OFT compared to males (*p*=0.011). **(C)** Representative track map for female **(C1)** control, **(C2)** eDREADD, and **(C3)** iDREADD mice. Blue and red circles denote the start and end of the track path, respectively. **(D)** There was no difference in the total distance traveled during the OFT. **(E)** There was no difference in the total distance traveled during the OFT in female and male cohorts.

Locomotor activity was also monitored (Figure 8C-E). Representative track maps are shown for female mice (8C1-C3). Note that some of the female eDREADD mice showed higher activity both within the center and periphery of the open field (Figure 8C2) but others did not, and there were no significant differences between the treatments. Quantification in Figure 8D-E was based on total distance traveled in the OFT and was analyzed by two-way ANOVA. There was no effect of treatment (*F*(2,67)=2.657, *p*=0.077; Figure 8D) or sex (*F*(1,67)=0.002, *p*=0.963; Figure 8E) on distance traveled in the OFT.

In summary, there was no significant effect of treatment, but a main effect of sex. Male mice, regardless of treatment showed similar behaviors, whereas female mice typically spent less time in the center of the OFT. This observation is consistent with sex differences in basal anxiety and exploration and can make interpretations of the OFT data challenging.

#### 3.1.8 EPM

Next, we evaluated anxiety-like behavior in the EPM. A two-way ANOVA revealed a significant effect of treatment (*F*(2,67)=3.379, *p*=0.040) but not sex (*F*(1,67)=0.299, *p*=0.586). Tukey’s post-hoc test showed that eDREADD mice spent a greater percent of time in the open arms (23.72 ± 3.64 %) compared to control mice (14.12 ± 0.89 %; *p*=0.033; Figure 9A). Other post-hoc comparisons were not significant (all *p* values >0.289). When data were separated so female and male cohorts could be compared, there were no significant effects of treatment or sex (Figure 9B). The lack of effect of treatment is consistent with a relatively small effect of eDREADD treatment in the pooled data (Figure 9A).

**Figure 9.**
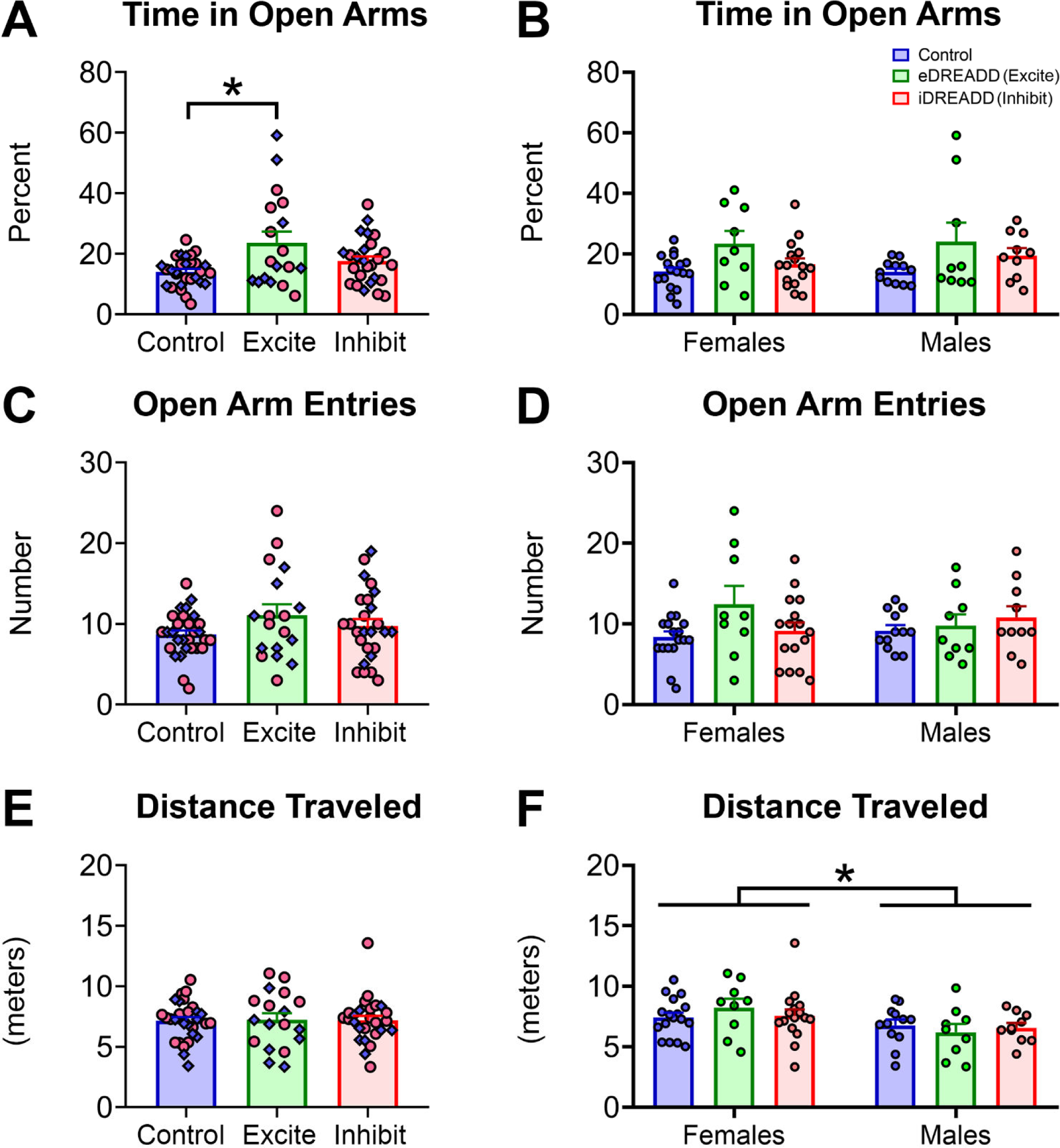
EPM in control, eDREADD and iDREADD mice. **(A)** eDREADD mice spent a greater percent of time in the open arms of the EPM compared to control mice (*p*=0.033). **(B)** There was no effect of treatment on the percent of time spent in the open arms of the EPM when pooled data in B were separated according to sex. **(C)** There was no effect of treatment on the number of open arm entries. **(D)** There was no effect of treatment on the number of open arm entries when pooled data in D were separated by sex. **(E)** There was no effect of treatment on the distance traveled during the EPM. **(F)** Female mice traveled a significantly greater distance than male mice during the EPM test (*p*=0.008). However, there was no effect of treatment in female and male cohorts.

The total number of open arm entries was also evaluated, and a two-way ANOVA found no effect of treatment (*F*(2,67)=0.723, *p*=0.488) or sex (*F*(1,67)=0.333, *p*=0.565; Figure 9C-D). Locomotor activity in the EPM was also evaluated by tracking the distance traveled during the test. A two-way ANOVA found no overall effect of treatment (*F*(2,67)=0.034, *p*=0.965; Figure 9E), but a significant effect of sex (*F*(1,67)=7.473, *p*=0.008), attributable to female mice (7.652 ± 0.299 meters) traveling a greater distance than male mice (6.547 ± 0.292 meters; Figure 9F). Notably, these results are consistent with sex differences in EPM behaviors (Belviranli et al., 2012; Scholl et al., 2019).

In summary, the results of the EPM suggest that eDREADD mice showed a modest increase in the time spent in the open arms of the EPM. Consistent with this small increase, there were no treatment differences in female or male cohorts. More time spent in the open arms is often interpreted as anxiolytic, but the small treatment effect suggest conclusions should be made with caution. Also, female mice traveled a greater distance than male mice and this result also suggests the EPM data should be cautiously interpreted.

### 3.2 MC effects on the DG circuit: c-Fos immunohistochemistry

C-fos immunoreactivity was used to confirm that MC activity was increased by eDREADD treatment and address whether iDREADD treatment reduced MC activity. Examining c-fos immunoreactivity after HCNOE was chosen because we have previously reported that the HCNOE task induces expression of c-Fos protein in a subset of MCs (Bernstein et al., 2019).

Therefore, mice were sacrificed 90 minutes after HCNOE to evaluate c-Fos protein in MCs. GCs were also examined to gain insight into potential effects of altered MC activity on GCs. Brains were cut in the coronal and horizontal plane to best evaluate dorsal and ventral hippocampus, as described in the Methods.

#### 3.2.1 Hilar c-Fos

First, c-Fos was analyzed in the hilus of coronal sections (as described in the Methods; Figure 10A). A two-way ANOVA revealed an effect of treatment (*F*(2,50)=80.42, *p*<0.001) and no effect of septotemporal location (*F*(1,50)=1.505, *p*=0.225). Tukey’s post-hoc test revealed that eDREADD mice (18.34 ± 2.17 cells) had a significantly greater average number of hilar c-Fos-immunoreactive cells compared to control (2.26 ± 0.20 cells) and iDREADD mice (2.32 ± 0.40 cells; all *p* values <0.001; Figure 10B). These findings are an important confirmation that eDREADD treatment increased neuronal activity of hilar neurons during this task. The hilar neurons were probably MCs because we previously found that HCNOE preferentially activates MCs compared to other hilar neurons after HCNOE (Duffy et al., 2013; Moretto et al., 2017; Bernstein et al., 2019) and DREADDs were preferentially expressed in MCs (Figure 1).

**Figure 10.**
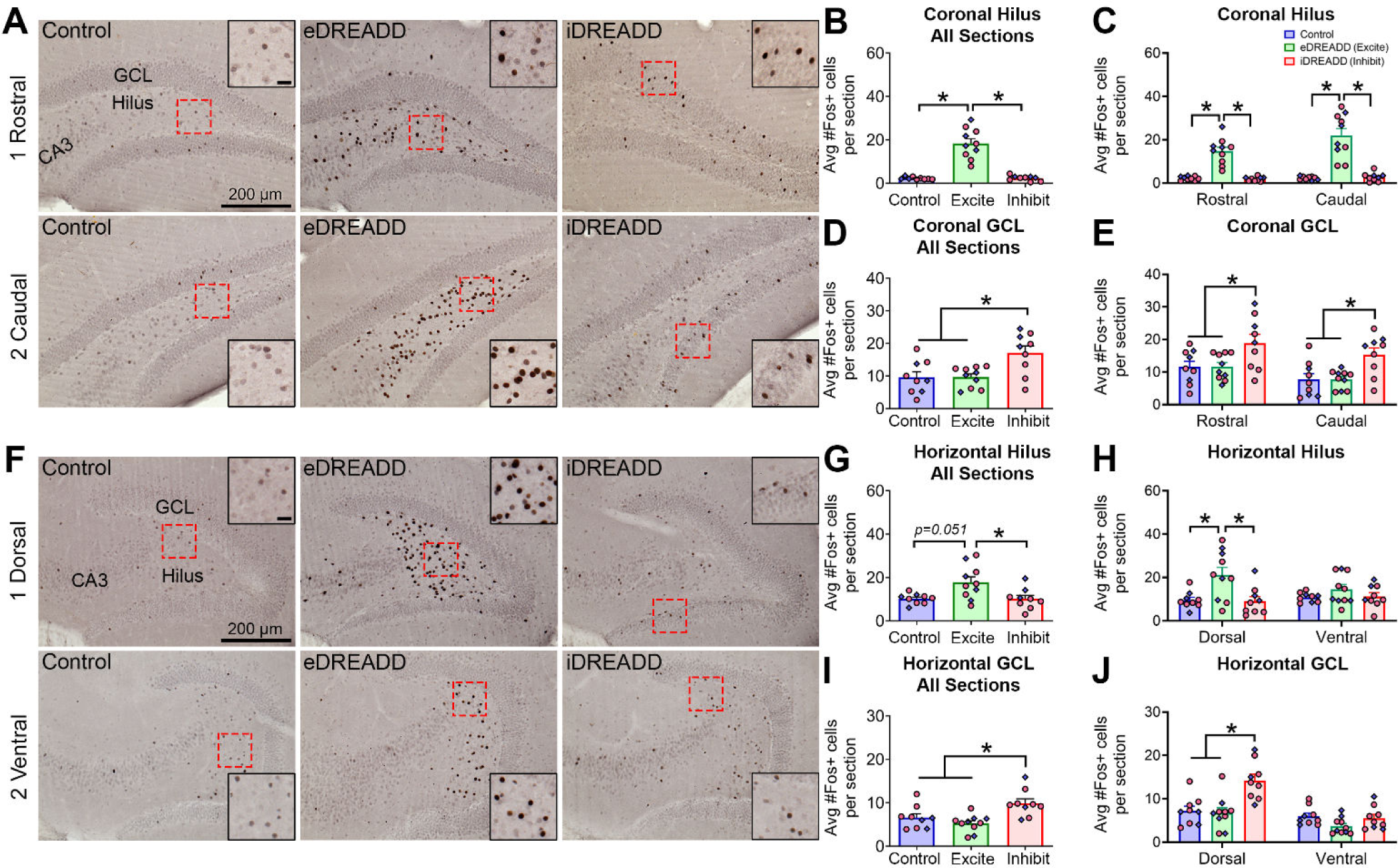
DREADD effects on hilar and GC c-Fos immunoreactivity. **(A)** 1-2. Representative c-Fos-immunoreactive (ir) in rostral and caudal coronal sections. Inset scalebar: 20µm. Control, eDREADD, and iDREADD mice were sacrificed 90 minutes after completing HCNOE to evaluate c-Fos-ir cells. Mice were treated with CNO 30 minutes before HCNOE. **(B)** All coronal sections of eDREADD mice had significantly more hilar c-Fos-ir cells per section (red arrows) than control and iDREADD mice (all *p* values <0.001). **(C)** When coronal sections were divided into rostral and caudal levels, both rostral and caudal sections had significantly more hilar c-Fos-ir cells per section in eDREADD mice compared to control and iDREADD mice (all *p* values <0.001). **(D)** All coronal sections of iDREADD mice had significantly more GCL c-Fos-ir cells per section than control and eDREADD mice (all *p* values <0.001). **(E)** When divided into rostral and caudal levels, both levels had significantly more GCL c-Fos-ir cells per section in iDREADD mice compared to control and eDREADD mice (all *p* values <0.017). **(F)** 1-2. Representative c-Fos-ir in dorsal and ventral horizontal sections. Inset scalebar: 20µm. **(G)** All horizontal sections of eDREADD mice had significantly more hilar c-Fos-ir cells per section than iDREADD mice (*p*=0.007). **(H)** In dorsal horizontal sections, eDREADD mice had significantly more hilar c-Fos-ir cells per section than control and iDREADD mice (all *p* values <0.028). There was no treatment difference in ventral sections. **(I)** All horizontal sections of iDREADD mice had significantly more GCL c-Fos-ir cells per section than control and eDREADD mice (all *p* values <0.007). **(J)** In dorsal horizontal sections, iDREADD mice had significantly more GCL c-Fos-ir cells per section than control and iDREADD mice (all *p* values <0.001). There was no treatment difference in ventral horizontal sections.

We also found that iDREADD treatment resulted in low levels of c-Fos immunoreactivity in the hilus. The controls also had a low level of hilar c-Fos, so the iDREADD-treated mice did not differ from controls. However, our prior studies of iDREADDs on patched MCs (using similar methods to what were used here) showed that CNO hyperpolarizes and reduces firing of MCs (Botterill et al., 2019). Therefore, it is likely that iDREADDs inhibited MCs, but due to the low c-Fos levels in control mice, it was difficult to detect a further reduction after iDREADD treatment. The low number of c-Fos-immunoreactive MCs in dorsal DG is consistent with prior studies of HCNOE (Bernstein et al., 2019; see also Duffy et al., 2013; Moretto et al., 2017)

Next, we compared relatively rostral and more caudal coronal sections. Tukey’s post-hoc tests revealed that in rostral sections, eDREADD mice (14.72 ± 1.87 cells) had significantly more hilar c-Fos cells per section than control (2.22 ± 0.31 cells) and iDREADD mice (1.90 ± 0.32 cells; all *p* values <0.001; Figure 10C). A similar result was observed in caudal sections, with more hilar c-Fos cells per section in eDREADD mice (21.96 ± 3.17 cells) compared to control (2.30 ± 0.28 cells) and iDREADD mice (2.73 ± 0.65 cells; all *p* values <0.001; Figure 10C).

Next we analyzed horizontal sections (Figure 10F). Sections were selected from relatively dorsal and ventral levels. A two-way ANOVA revealed an effect of treatment (*F*(2,50)=5.540, *p*<0.001) and no effect of septotemporal location (*F*(1,50)=0.121, *p*=0.728). Tukey’s post-hoc test revealed that the average number of hilar c-Fos-immunoreactive cells was greater in eDREADD mice (17.83 ± 2.45 cells) compared to iDREADD mice (10.17 ± 1.57 cells; *p*=0.007; Figure 10G). Control mice (10.20 ± 0.78 cells) did not differ from either treatment (all *p* values >0.051). Tukey’s post-hoc test further revealed that in dorsal horizontal sections, eDREADD mice (21.15 ± 3.53 cells) had a greater number of c-Fos-immunoreactive cells per section than control (9.56 ± 1.30 cells) and iDREADD mice (9.08 ± 2.16 cells; all *p* values <0.028; Figure 10H).

There were no differences between eDREADD, iDREADD and control mice in the numbers of hilar c-Fos-immunoreactive cells per section in ventral horizontal sections (all *p* values >0.529). The results are likely to be related to the viral injection sites, which were probably did not reach the most ventral part of the DG (see Methods). Although Figure 1 shows fairly strong expression in dorsal and caudal coronal sections, the extreme temporal (ventral) pole showed few MC somata expressing mCherry.

In summary, eDREADD treatment increased hilar c-Fos-immunoreactive cells in a robust manner, except for the most ventral part of the DG which may have had less somatic expression of DREADDs. iDREADD treatment did not significantly decrease hilar c-Fos immunoreactivity, which could be due to low numbers of c-Fos cells in controls.

#### 3.2.2 GCL c-Fos

Next, we evaluated c-Fos in the GCL to gain insight into whether MC excitation or inhibition influenced the activity of GCs. Past studies found that the vast majority of c-Fos-immunoreactive cells in the GCL after exploration of novel objects express markers of GCs rather than GABAergic neurons (Duffy et al., 2013; Bernstein et al., 2019), so we infer c-Fos-immunoreactive cells in the GCL were GCs below. Notably, GABAergic neurons do not appear to express c-Fos readily after these behaviors (Duffy et al., 2013; Moretto et al., 2017; Bernstein et al., 2019), limiting what can be concluded about their roles.

A two-way ANOVA revealed a significant effect of treatment (*F*(2,50)=11.24, *p*<0.001). Tukey’s post-hoc test indicated that iDREADD mice (17.07 ± 2.13 cells) had a greater average number of c-Fos-immunoreactive cells in the GCL compared to control (9.60 ± 1.68 cells) and eDREADD mice (9.67 ± 0.98 cells; all *p* values <0.001; Figure 10D). This result suggests that GCs are activated by iDREADD treatment. One explanation is that iDREADD treatment reduces the activity in the indirect MC→GABAergic neuron→GC pathway, resulting in a net increase in GC activation, which is a hypothesis supported by prior studies that suggest MC loss promotes GC excitability (Sloviter, 1991; Jinde et al., 2012).

We also observed a main effect of septotemporal location (*F*(1,50)=6.66, *p*=0.012) on coronal GCL c-Fos immunoreactivity. This effect was attributable to rostral sections having greater c-Fos immunoreactivity than caudal sections (Figure 10E), consistent with past studies (Bernstein et al., 2019). In rostral coronal sections, Tukey’s post-hoc tests found that number of c-Fos cells in the GCL was greater in iDREADD mice (18.85 ± 2.69 cells) compared to control (11.54 ± 1.74 cells) and eDREADD mice (11.65 ± 1.25 cells; all *p* values <0.017; Figure 10E). Similarly, in caudal coronal sections, the number of c-Fos cells in the GCL was significantly greater in iDREADD mice (15.30 ± 2.06 cells) compared to control (7.67 ± 1.80 cells) and eDREADD mice (7.71 ± 0.89 cells; all *p* values <0.013; Figure 10E).

We also evaluated the number of c-Fos cells in the GCL of horizontal sections. A two-way ANOVA revealed significant effect of treatment (*F*(2,50)=10.91, *p*<0.001), septotemporal location (*F*(1,50)=26.90, *p*<0.001), and a significant interaction (*F*(2,50)=7.112, *p*=0.001). Tukey’s post-hoc tests showed that iDREADD mice had a greater number of c-Fos immunoreactive cells in the GCL (9.87 ± 1.01 cells) compared to control (6.59 ± 0.88 cells) and eDREADD mice (5.24 ± 0.62 cells; all *p* values <0.031; Figure 10I). For dorsal horizontal sections, the average number of c-Fos cells in the GCL was significantly greater in iDREADD mice (14.22 ± 1.44 cells) compared to control (7.20 ± 1.09 cells) and eDREADD mice (6.76 ± 1.17 cells; all *p* values <0.001; Figure 10J). In the most ventral horizontal sections, there were no significant differences between eDREADD, iDREADD and control mice (all *p* values >0.264).

In summary, the results show contrasting effects of DREADDs on the DG circuit. The MC c-Fos data suggest that eDREADDs significantly increased MC activity as one would predict, given the excitatory actions of eDREADDs. However, iDREADDs did not have the opposite effect, presumably due to the low levels of MC c-Fos in control mice.

Regarding GC c-Fos, the results can be explained by the two circuits that MCs use to influence GCs: the direct MC-GC pathway which excites GCs and the indirect MC→GABAergic neuron→GC pathway which inhibits GCs (Figure 1E). The indirect pathway appears to dominate under standard conditions (Jinde et al., 2012; Hsu et al., 2016; Bui et al., 2018; Yeh et al., 2018). After eDREADD activation by CNO, there would be greater activation of both the direct and indirect pathways which appeared to cause no net change in GC c-Fos (Figure 11A). In contrast, iDREADD inhibition of MCs might be effective in reducing the indirect pathway and disinhibit GCs (Figure 11B). Then when an animal is exposed to novel objects, excitatory input from entorhinal cortex (carrying spatial and object information; Eichenbaum et al., 2012; Knierim et al., 2014; Knierim and Neunuebel, 2016) would be much more likely to cause GC firing.

**Figure 11.**
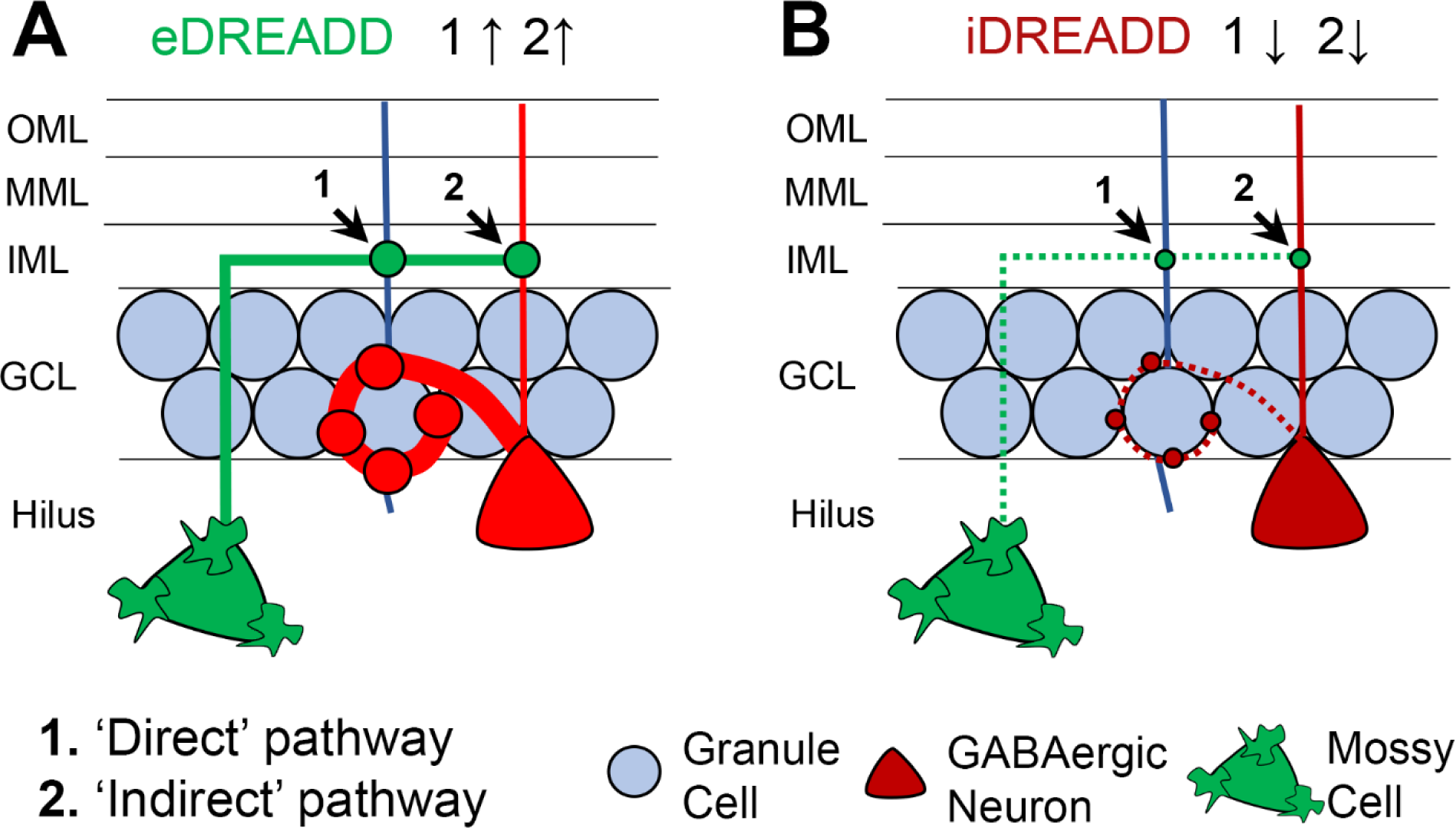
DREADD effects on the MC circuit. **(A)** eDREADD treatment increases MC firing and neurotransmitter release, which would facilitate both the *(1)* direct MC→GC and *(2)* indirect MC→GABAergic neuron→GC pathways. Notably, eDREADD-treatment had a minimal effect on GCL c-Fos-ir, possibly due to opposing excitatory and inhibitory effects at the direct and indirect pathways, respectively. **(B)** iDREADD treatment inhibits MC firing and neurotransmitter release, which would reduce MC effects at the *(1)* direct MC→GC and *(2)* indirect MC→GABAergic neuron→GC pathways. The reduced drive at the direct and indirect pathways appeared to promote GC firing, since iDREADD-treated mice showed significantly greater GCL c-Fos immunoreactivity. This finding is supported by previous studies that suggest that MC loss promotes GC excitability (Sloviter, 1991; Jinde et al., 2012 but see Ratzliff et al., 2004).

Taken together, the results of eDREADD and iDREADD treatment are consistent with a relative dominance of the indirect pathway under standard conditions (Figure 11). If one now turns to the implications for behavior, the c-Fos results suggest that increased MC activity by eDREADDs may cause competing effects on the direct and indirect pathways. If the indirect pathway is normally dominant, GCs would be more inhibited. That effect appears to worsen some anxiety-like behaviors and cognitive tasks. Conversely, inhibition of MCs would lead to more activity of GCs if the indirect pathway is dominant. That effect appeared to lessen some of the anxiety-like behaviors and improve some of the cognitive tasks. The implication is that more GC activity improves some types of behavior, consistent with increased GC firing allowing a greater DG influence in the networks regulating behavior. Another possibility is that increased GC activity promotes GC expression of activity-dependent transcription factors underlying synaptic plasticity, and greater encoding of experience within the DG.

## 4. DISCUSSION

The present study examined the role of MCs in cognitive and anxiety-like behaviors using a gain- and loss-of function approach. Remarkably, exciting versus inhibiting MCs produced opposing behavioral effects in several tasks (e.g., CFC, NOR, HCNOE, NSF). Exciting or inhibiting MCs also resulted in behaviors that were significantly different from control mice in several tasks (e.g., CFC, NOR, HCNOE, NSF, LDB, EPM). These results support the hypothesis that MCs influence cognitive and anxiety-like behaviors in mice.

### 4.1 MCs influence cognitive behaviors

There has been a lot of work to understand the role of MCs in DG functions related to spatial navigation, spatial memory and a widely discussed function of the DG known as pattern separation (Danielson et al., 2017; GoodSmith et al., 2017; Senzai and Buzsaki, 2017; GoodSmith et al., 2019; Jung et al., 2019). Past studies have also addressed how MCs and GCs interact with area CA3 to support these functions (Penttonen et al., 1997; Lisman, 1999; Scharfman, 2007a; Myers and Scharfman, 2009; Myers and Scharfman, 2011; Knierim and Neunuebel, 2016; GoodSmith et al., 2019). There also are a number of studies which addressed the role of MCs in functions of the DG related to novelty, both novelty in location and object novelty (Jinde et al., 2012; Duffy et al., 2013; Moretto et al., 2017; Bernstein et al., 2019) but methods involved neuronal damage to MCs, or only used anatomical methods.

Therefore, the results are timely. For the tests we discuss as ‘cognitive’, we investigated contextual memory (CFC) and novel object tests (NOR, NOL, HCNOE). The results show that exciting MCs with eDREADDs significantly impaired contextual fear learning and memory. Our finding contrasts with Jinde and colleagues (Jinde et al., 2012) who reported that ablation of MCs impaired contextual discrimination. However, Jinde et al. used a different task and reduced MC activity through MC ablation which are likely to cause complex secondary changes.

We found few effects in NOL but robust effects in NOR and HCNOE. In NOL, exciting or inhibiting MCs had no significant effects on the training or testing DI in the NOL task. In contrast, exciting MCs significantly impaired the NOR testing DI without an effect on the training DI. Our results differ from (Bui et al., 2018), who reported that MC photoinhibition during the learning phase of an object location task impaired location memory, without an effect on object recognition learning and memory. Notably, methodological differences may account for the discrepancies. For example, Bui and colleagues moved the object location approximately half the distance as in the present study, which is notable because it has been reported that the DG is critical for small but not large spatial discrimination (Clelland et al., 2009; Schmidt et al., 2012). This idea is supported by a recent optogenetic study that reported MCs were sensitive to small but not large spatial displacement in a touchscreen task (Jung et al., 2019). Our results also differ from Bui et al. (Bui et al., 2018) in that their training and testing interval was 24 hours and photoinhibition was sensitive to learning. In contrast, our training and testing interval was one hour apart and CNO was injected before training. The effects of DREADDs after CNO injection are known to last for several hours (Smith et al., 2016), and therefore our approach provided sustained DREADD effects.

In HCNOE, inhibiting and exciting MCs resulted in the highest and lowest levels of object exploration, respectively. These results support the view that eDREADDs interfere with processing information about novelty, whereas iDREADDs facilitate exploration. If MCs excite the circuit too much or for too long, adverse effects would seem likely, as recent study demonstrates (Botterill et al., 2019). If iDREADDs are anxiolytic, then it seems reasonable that animals would explore more.

#### 4.2.1 MCs influence anxiety-like behaviors

There is good reason to examine MCs in anxiety-like behavior. One reason is the DG appears to regulate the response to behavioral stress and associated anxiety-like behavior, especially the ventral DG (Anacker et al., 2018). Importantly, MCs in dorsal DG project to ventral DG (Scharfman, 2016). Also, MCs appear to have a role in depression (Oh et al., 2019) which usually occurs with anxiety. MCs express genes that are linked to schizophrenia (Scharfman and Bernstein, 2015; Yuan et al., 2015), which is a disease with anxiety (Temmingh and Stein, 2015). In addition, the DG is influenced by behavioral stress (McEwen et al., 2016), which often leads to anxiety, and stress can reduce c-Fos in MCs (Moretto et al., 2017).

#### 4.2.2 MCs have a role in anxiety

Although there have been several studies about the role of MCs in functions of the DG related to cognition (see Section 4.1), fewer studies have addressed the role of MCs in anxiety-like behavior. Also, few studies have examined both anxiety-like behavior and cognition in the same study. Therefore, our results led to some significant insights.

First, the results suggest that MCs have a role in anxiety-like behavior, but it appears to be selective. This notion is consistent with DG functions, which are critical only to some types of anxiety-like behavior. DREADD effects were found in tasks that are commonly used to probe anxiety (NSF, LDB, EPM) except OFT. Notably, a recent study also reported trends but no significant effects of DREADDs on MCs in OFT (Oh et al., 2019). However, Jinde and colleagues reported that ablation of MCs resulted in anxiety-like phenotypes in the OFT (Jinde et al., 2012), but there were methodological limitations as described above.

In many tasks we tested, iDREADDs were anxiolytic but eDREADDs were anxiolytic in the EPM. A similar anxiolytic effect of MC excitation in the EPM was recently reported (Oh et al., 2019). In contrast, (Bui et al., 2018) found no effect of MC inhibition in the EPM, but their methods were much different.

Taken together, tasks that involved animals moving into a large open field or elevated area without objects (OFT, EPM) seemed to show different results from tasks that involved a smaller area (LDB, HCNOE), or involved objects (NSF, HCNOE). Therefore, the context of a large open space may influence when MCs are involved. The importance of objects is consistent with the role of the DG in differentiating contexts in CFC but not cued conditioning (Phillips and LeDoux, 1992).

#### 4.2.3 The role of MCs in anxiety could regulate cognitive performance

The results suggest a hypothesis: the role of MCs in cognition could be related to the MC role in anxiety-like behavior. This hypothesis is suggested by the data showing that iDREADDs often decreased anxiety-like behavior, and iDREADDs also improved performance on some cognitive tasks. Conversely, eDREADDs often worsened cognitive tasks. It is intriguing to consider that cognitive functions of the DG could be gated by the degree of anxiety, and the gate could involve MCs.

### 4.3 Roles of MC and GC activity in behavior

A common question is how DG circuitry is involved in anxiety-like and cognitive behavior. Past studies and the c-Fos data presented here provide a working hypothesis. Thus, two pathways have been proposed to explain MC effects on GCs, the direct excitatory MC→GC pathway and the indirect inhibitory MC→GABAergic neuron→GC pathway (Figure 1E). Prior work suggests a relative dominance of the indirect pathway over the direct pathway under standard conditions (Sloviter, 1991; Jinde et al., 2012; Hsu et al., 2016; Bui et al., 2018; Yeh et al., 2018). Our data showing that eDREADDs led to little effect on GC c-Fos suggests that increasing the already strong inhibition of GCs did not have much effect (Figure 11A). However, eDREADDs did have adverse effects behaviorally, presumably because synchronous, sustained activation of the majority of MCs is nonphysiological and therefore disrupts normal DG function.

Use of iDREADDs to inhibit a large number of MCs led to a robust excitatory effect on GC c-Fos, suggesting iDREADDs reduced the indirect inhibitory pathway and this led to GC excitation (Figure 11B). Here the behavioral effect was positive, possibly because the E:I balance of GCs is normally biased toward inhibition, and for optimal behavior a little more GC activity is beneficial.

### 4.4 Sex-dependent behavioral effects

The majority of studies to date on MCs have focused on male subjects, which is problematic because females and males have different basal anxiety-like behavior and often utilize different cognitive strategies than male subjects (Galea et al., 2017). Examples of female-specific effects include fear learning and memory in the CFC, more robust exploration in the HCNOE task, latency to feed in the NSF, time in the center of the OFT, and distance traveled in the EPM. These data suggest that previous studies, which typically used males only, might have underestimated behavioral effects of MCs by focusing on male subjects alone.

There are reasons why some effects might have been more robust in females. For example, estrogen increases the neurotrophin brain-derived neurotrophic factor (BDNF) in GCs, which is important to the DG because BDNF regulates DG structure, function, and plasticity (Harte-Hargrove et al., 2013; Scharfman and MacLusky, 2014). Higher BDNF protein in GC axons (mossy fibers) increases activation of CA3 by GCs and improves NOL performance (Scharfman et al., 2003; Scharfman, 2007b; Skucas et al., 2013). BDNF is particularly relevant to MCs because MCs exhibit a BDNF-dependent form of long-term potentiation specifically at MC→GC synapses (Hashimotodani et al., 2017).

### 4.5 Implications for disease

One of the central hypotheses about MCs in disease is about temporal lobe epilepsy (TLE), where it has been shown that substantial loss of MCs occurs (Scharfman, 2016). Removal of MCs from the circuitry has been suggested to promote epilepsy because there is reduced activity of the MC→GABAergic neuron→GC pathway (Sloviter et al., 2003). As a result, GCs become hyperexcitable and lead to hyperexcitability in hippocampus. Support for this hypothesis, and alternative hypotheses, have been presented intermittently since the 1990’s (Sloviter et al., 2003; Ratzliff et al., 2004; Jinde et al., 2012; Scharfman, 2016; Bui et al., 2018). In contrast to the view that MC loss has adverse effects in TLE, the data provided here suggest this is not true in the normal brain, where inhibiting MCs had some beneficial effects and exciting MCs has some adverse effects. The different roles of MCs in disease compared to normal conditions might be due to large changes in the DG in TLE (de Lanerolle et al., 2012; Dingledine et al., 2017; Danzer, 2018), but it is also possible that the role of MCs changes radically, depending on the behavior.

## 5 CONCLUSIONS

Here, we used a gain- and loss-of function approach to study MCs in cognitive and affective behaviors in female and male mice. Manipulations of MCs led to altered behavioral responses in numerous cognitive and anxiety-like behaviors. Furthermore, exciting vs inhibiting MCs led to distinct patterns of hilar and GC c-Fos immunoreactivity, indicating that MC activity influences the DG. Collectively, this study provides evidence that MCs influence cognitive and anxiety-like behaviors in male and female mice.

## 6 DATA AVAILABILITY STATEMENT

The datasets generated for this study are available upon request to the corresponding author.

## 8 AUTHOR CONTRIBUTIONS

Conceptualization: JJB, KYV, HES. Data collection and analysis: JJB, KYV, KJG, CMT, JJL.

Wrote the manuscript: JJB, KYV, HES. All authors reviewed and approved the manuscript.

## REFERENCES

Altemus M, Sarvaiya N, Neill Epperson C (2014) Sex differences in anxiety and depression clinical perspectives. Front Neuroendocrinol 35:320–330.

Anacker C, Hen R (2017) Adult hippocampal neurogenesis and cognitive flexibility - linking memory and mood. Nat Rev Neurosci 18:335–346.

Anacker C, Luna VM, Stevens GS, Millette A, Shores R, Jimenez JC, Chen B, Hen R (2018) Hippocampal neurogenesis confers stress resilience by inhibiting the ventral dentate gyrus. Nature 559:98–102.

Azevedo EP, Pomeranz L, Cheng J, Schneeberger M, Vaughan R, Stern SA, Tan B, Doerig K, Greengard P, Friedman JM (2019) A role of Drd2 hippocampal neurons in context-dependent food intake. Neuron 102:873–886 e875.

Bailey KR, Crawley JN (2009) Anxiety-related behaviors in mice. In: Methods of Behavior Analysis in Neuroscience (nd, Buccafusco JJ, eds). Boca Raton (FL).

Bale TL, Epperson CN (2015) Sex differences and stress across the lifespan. Nat Neurosci 18:1413–1420.

Belviranli M, Atalik KE, Okudan N, Gokbel H (2012) Age and sex affect spatial and emotional behaviors in rats: the role of repeated elevated plus maze test. Neuroscience 227:1–9.

Bernstein HL, Lu YL, Botterill JJ, Scharfman HE (2019) Novelty and novel objects increase c-Fos immunoreactivity in mossy cells in the mouse dentate gyrus. Neural Plast 2019:1815371.

Bernstein HL, Lu YL, Botterill JJ, Duffy AM, LaFrancois J, Scharfman H (2020) Excitatory effects of dentate gyrus mossy cells and their ability to influence granule cell firing: an optogenetic study in adult mouse hippocampal slices. bioRxiv https://doiorg/101101/20200606137844.

Botterill JJ, Brymer KJ, Caruncho HJ, Kalynchuk LE (2015a) Aberrant hippocampal neurogenesis after limbic kindling: Relationship to BDNF and hippocampal-dependent memory. Epilepsy Behav 47:83–92.

Botterill JJ, Guskjolen AJ, Marks WN, Caruncho HJ, Kalynchuk LE (2015b) Limbic but not non-limbic kindling impairs conditioned fear and promotes plasticity of NPY and its Y2 receptor. Brain Struct Funct 220:3641–3655.

Botterill JJ, Fournier NM, Guskjolen AJ, Lussier AL, Marks WN, Kalynchuk LE (2014) Amygdala kindling disrupts trace and delay fear conditioning with parallel changes in Fos protein expression throughout the limbic brain. Neuroscience 265:158–171.

Botterill JJ, Lu YL, LaFrancois JJ, Bernstein HL, Alcantara-Gonzalez D, Jain S, Leary P, Scharfman HE (2019) An excitatory and epileptogenic effect of dentate gyrus mossy cells in a mouse model of epilepsy. Cell Rep 29:2875–2889 e2876.

Brymer KJ, Johnston J, Botterill JJ, Romay-Tallon R, Mitchell MA, Allen J, Pinna G, Caruncho HJ, Kalynchuk LE (2020) Fast-acting antidepressant-like effects of Reelin evaluated in the repeated- corticosterone chronic stress paradigm. Neuropsychopharmacology.

Bui AD, Nguyen TM, Limouse C, Kim HK, Szabo GG, Felong S, Maroso M, Soltesz I (2018) Dentate gyrus mossy cells control spontaneous convulsive seizures and spatial memory. Science 359:787–790.

Clelland CD, Choi M, Romberg C, Clemenson GD, Jr., Fragniere A, Tyers P, Jessberger S, Saksida LM, Barker RA, Gage FH, Bussey TJ (2009) A functional role for adult hippocampal neurogenesis in spatial pattern separation. Science 325:210–213.

Danielson NB, Turi GF, Ladow M, Chavlis S, Petrantonakis PC, Poirazi P, Losonczy A (2017) In vivo imaging of dentate gyrus mossy cells in behaving mice. Neuron 93:552–559 e554.

Danzer SC (2018) Contributions of adult-generated granule cells to hippocampal pathology in temporal lobe epilepsy: a neuronal bestiary. Brain Plast 3:169–181.

de Lanerolle NC, Lee TS, Spencer DD (2012) Histopathology of human epilepsy. In: Jasper’s Basic Mechanisms of the Epilepsies (th, Noebels JL, Avoli M, Rogawski MA, Olsen RW, Delgado-Escueta AV, eds). Bethesda (MD).

Demireva EY, Suri D, Morelli E, Mahadevia D, Chuhma N, Teixeira CM, Ziolkowski A, Hersh M, Fifer J, Bagchi S, Chemiakine A, Moore H, Gingrich JA, Balsam P, Rayport S, Ansorge MS (2018) 5- HT2C receptor blockade reverses SSRI-associated basal ganglia dysfunction and potentiates therapeutic efficacy. Mol Psychiatry.

Dingledine R, Coulter DA, Fritsch B, Gorter JA, Lelutiu N, McNamara J, Nadler JV, Pitkanen A, Rogawski MA, Skene P, Sloviter RS, Wang Y, Wadman WJ, Wasterlain C, Roopra A (2017) Transcriptional profile of hippocampal dentate granule cells in four rat epilepsy models. Sci Data 4:170061.

Donner NC, Lowry CA (2013) Sex differences in anxiety and emotional behavior. Pflugers Arch 465:601–626.

Duffy AM, Schaner MJ, Chin J, Scharfman HE (2013) Expression of c-fos in hilar mossy cells of the dentate gyrus in vivo. Hippocampus 23:649–655.

Dulawa SC, Hen R (2005) Recent advances in animal models of chronic antidepressant effects: the novelty-induced hypophagia test. Neurosci Biobehav Rev 29:771–783.

Eichenbaum H, Sauvage M, Fortin N, Komorowski R, Lipton P (2012) Towards a functional organization of episodic memory in the medial temporal lobe. Neurosci Biobehav Rev 36:1597–1608.

Fanselow MS (1980) Conditioned and unconditional components of post-shock freezing. Pavlov J Biol Sci 15:177–182.

Fanselow MS, Pennington ZT (2017) The danger of LeDoux and Pine’s two-system framework for fear. Am J Psychiatry 174:1120–1121.

Galea LAM, Frick KM, Hampson E, Sohrabji F, Choleris E (2017) Why estrogens matter for behavior and brain health. Neurosci Biobehav Rev 76:363–379.

GoodSmith D, Lee H, Neunuebel JP, Song H, Knierim JJ (2019) Dentate gyrus mossy cells share a role in pattern separation with dentate granule cells and proximal CA3 pyramidal cells. J Neurosci 39:9570–9584.

GoodSmith D, Chen X, Wang C, Kim SH, Song H, Burgalossi A, Christian KM, Knierim JJ (2017) Spatial representations of granule cells and mossy cells of the dentate gyrus. Neuron 93:677–690 e675.

Guidi S, Severi S, Ciani E, Bartesaghi R (2006) Sex differences in the hilar mossy cells of the guinea- pig before puberty. Neuroscience 139:565–576.

Guskjolen A, Kenney JW, de la Parra J, Yeung BA, Josselyn SA, Frankland PW (2018) Recovery of “lost” infant memories in mice. Curr Biol 28:2283–2290 e2283.

Hajszan T, Milner TA, Leranth C (2007) Sex steroids and the dentate gyrus. Prog Brain Res 163:399–415.

Harte-Hargrove LC, Maclusky NJ, Scharfman HE (2013) Brain-derived neurotrophic factor-estrogen interactions in the hippocampal mossy fiber pathway: implications for normal brain function and disease. Neuroscience 239:46–66.

Harte-Hargrove LC, Varga-Wesson A, Duffy AM, Milner TA, Scharfman HE (2015) Opioid receptor- dependent sex differences in synaptic plasticity in the hippocampal mossy fiber pathway of the adult rat. J Neurosci 35:1723–1738.

Hashimotodani Y, Nasrallah K, Jensen KR, Chavez AE, Carrera D, Castillo PE (2017) LTP at hilar mossy cell-dentate granule cell synapses modulates dentate gyrus output by increasing excitation/inhibition balance. Neuron 95:928–943 e923.

Hsu TT, Lee CT, Tai MH, Lien CC (2016) Differential recruitment of dentate gyrus interneuron types by commissural versus perforant pathways. Cereb Cortex 26:2715–2727.

Jinde S, Zsiros V, Nakazawa K (2013) Hilar mossy cell circuitry controlling dentate granule cell excitability. Front Neural Circuits 7:14.

Jinde S, Zsiros V, Jiang Z, Nakao K, Pickel J, Kohno K, Belforte JE, Nakazawa K (2012) Hilar mossy cell degeneration causes transient dentate granule cell hyperexcitability and impaired pattern separation. Neuron 76:1189–1200.

Jung D, Kim S, Sariev A, Sharif F, Kim D, Royer S (2019) Dentate granule and mossy cells exhibit distinct spatiotemporal responses to local change in a one-dimensional landscape of visual- tactile cues. Sci Rep 9:9545.

Keiser AA, Turnbull LM, Darian MA, Feldman DE, Song I, Tronson NC (2017) Sex differences in context fear generalization and recruitment of hippocampus and amygdala during retrieval. Neuropsychopharmacology 42:397–407.

Kesner RP (2018) An analysis of dentate gyrus function (an update). Behav Brain Res 354:84–91.

Kim WB, Cho JH (2020) Encoding of contextual fear memory in hippocampal-amygdala circuit. Nat Commun 11:1382.

Kinnavane L, Albasser MM, Aggleton JP (2015) Advances in the behavioural testing and network imaging of rodent recognition memory. Behav Brain Res 285:67–78.

Klemenhagen KC, Gordon JA, David DJ, Hen R, Gross CT (2006) Increased fear response to contextual cues in mice lacking the 5-HT1A receptor. Neuropsychopharmacology 31:101–111.

Knierim JJ, Neunuebel JP (2016) Tracking the flow of hippocampal computation: Pattern separation, pattern completion, and attractor dynamics. Neurobiol Learn Mem 129:38–49.

Knierim JJ, Neunuebel JP, Deshmukh SS (2014) Functional correlates of the lateral and medial entorhinal cortex: objects, path integration and local-global reference frames. Philos Trans R Soc Lond B Biol Sci 369:20130369.

Komada M, Takao K, Miyakawa T (2008) Elevated plus maze for mice. J Vis Exp.

LeDoux JE, Pine DS (2016) Using neuroscience to help understand fear and anxiety: A two-system framework. Am J Psychiatry 173:1083–1093.

Leger M, Quiedeville A, Bouet V, Haelewyn B, Boulouard M, Schumann-Bard P, Freret T (2013) Object recognition test in mice. Nat Protoc 8:2531–2537.

Lisman JE (1999) Relating hippocampal circuitry to function: recall of memory sequences by reciprocal dentate-CA3 interactions. Neuron 22:233–242.

Luine VN, Beck KD, Bowman RE, Frankfurt M, Maclusky NJ (2007) Chronic stress and neural function: accounting for sex and age. J Neuroendocrinol 19:743–751.

MacLaren DA, Browne RW, Shaw JK, Krishnan Radhakrishnan S, Khare P, Espana RA, Clark SD (2016) Clozapine N-oxide administration produces behavioral effects in Long-Evans rats: implications for designing DREADD experiments. eNeuro 3.

Manvich DF, Webster KA, Foster SL, Farrell MS, Ritchie JC, Porter JH, Weinshenker D (2018) The DREADD agonist clozapine N-oxide (CNO) is reverse-metabolized to clozapine and produces clozapine-like interoceptive stimulus effects in rats and mice. Sci Rep 8:3840.

McEwen BS, Nasca C, Gray JD (2016) Stress effects on neuronal structure: hippocampus, amygdala, and prefrontal cortex. Neuropsychopharmacology 41:3–23.

Moretto JN, Duffy AM, Scharfman HE (2017) Acute restraint stress decreases c-fos immunoreactivity in hilar mossy cells of the adult dentate gyrus. Brain Struct Funct 222:2405–2419.

Myers CE, Scharfman HE (2009) A role for hilar cells in pattern separation in the dentate gyrus: a computational approach. Hippocampus 19:321–337.

Myers CE, Scharfman HE (2011) Pattern separation in the dentate gyrus: a role for the CA3 backprojection. Hippocampus 21:1190–1215.

Oh SJ, Cheng J, Jang JH, Arace J, Jeong M, Shin CH, Park J, Jin J, Greengard P, Oh YS (2019) Hippocampal mossy cell involvement in behavioral and neurogenic responses to chronic antidepressant treatment. Mol Psychiatry.

Palanza P (2001) Animal models of anxiety and depression: how are females different? Neurosci Biobehav Rev 25:219–233.

Penttonen M, Kamondi A, Sik A, Acsady L, Buzsaki G (1997) Feed-forward and feed-back activation of the dentate gyrus in vivo during dentate spikes and sharp wave bursts. Hippocampus 7:437–450.

Phillips RG, LeDoux JE (1992) Differential contribution of amygdala and hippocampus to cued and contextual fear conditioning. Behav Neurosci 106:274–285.

Ratzliff A, Santhakumar V, Howard A, Soltesz I (2002) Mossy cells in epilepsy: rigor mortis or vigor mortis? Trends Neurosci 25:140–144.

Ratzliff A, Howard AL, Santhakumar V, Osapay I, Soltesz I (2004) Rapid deletion of mossy cells does not result in a hyperexcitable dentate gyrus: implications for epileptogenesis. J Neurosci 24:2259–2269.

Scharfman HE (2007a) The CA3 “backprojection” to the dentate gyrus. Prog Brain Res 163:627–637.

Scharfman HE (2007b) The dentate gyrus: a comprehensive guide to structure, function, and clinical implications: Elsevier.

Scharfman HE (2016) The enigmatic mossy cell of the dentate gyrus. Nat Rev Neurosci 17:562–575.

Scharfman HE (2017) Advances in understanding hilar mossy cells of the dentate gyrus. Cell Tissue Res.

Scharfman HE, MacLusky NJ (2014) Differential regulation of BDNF, synaptic plasticity and sprouting in the hippocampal mossy fiber pathway of male and female rats. Neuropharmacology 76 Pt C:696–708.

Scharfman HE, Bernstein HL (2015) Potential implications of a monosynaptic pathway from mossy cells to adult-born granule cells of the dentate gyrus. Front Syst Neurosci 9:112.

Scharfman HE, Mercurio TC, Goodman JH, Wilson MA, MacLusky NJ (2003) Hippocampal excitability increases during the estrous cycle in the rat: a potential role for brain-derived neurotrophic factor. J Neurosci 23:11641–11652.

Schmidt B, Marrone DF, Markus EJ (2012) Disambiguating the similar: the dentate gyrus and pattern separation. Behav Brain Res 226:56–65.

Scholl JL, Afzal A, Fox LC, Watt MJ, Forster GL (2019) Sex differences in anxiety-like behaviors in rats. Physiol Behav 211:112670.

Seibenhener ML, Wooten MC (2015) Use of the open field maze to measure locomotor and anxiety-like behavior in mice. J Vis Exp:e52434.

Senzai Y, Buzsaki G (2017) Physiological properties and behavioral correlates of hippocampal granule cells and mossy cells. Neuron 93:691–704.

Simpson J, Kelly JP (2012) An investigation of whether there are sex differences in certain behavioural and neurochemical parameters in the rat. Behav Brain Res 229:289–300.

Skucas VA, Duffy AM, Harte-Hargrove LC, Magagna-Poveda A, Radman T, Chakraborty G, Schroeder CE, MacLusky NJ, Scharfman HE (2013) Testosterone depletion in adult male rats increases mossy fiber transmission, LTP, and sprouting in area CA3 of hippocampus. J Neurosci 33:2338–2355.

Sloviter RS (1991) Permanently altered hippocampal structure, excitability, and inhibition after experimental status epilepticus in the rat: the “dormant basket cell” hypothesis and its possible relevance to temporal lobe epilepsy. Hippocampus 1:41–66.

Sloviter RS, Zappone CA, Harvey BD, Bumanglag AV, Bender RA, Frotscher M (2003) “Dormant basket cell” hypothesis revisited: relative vulnerabilities of dentate gyrus mossy cells and inhibitory interneurons after hippocampal status epilepticus in the rat. J Comp Neurol 459:44–76.

Smith KS, Bucci DJ, Luikart BW, Mahler SV (2016) DREADDS: Use and application in behavioral neuroscience. Behav Neurosci 130:137–155.

Soltesz I, Bourassa J, Deschenes M (1993) The behavior of mossy cells of the rat dentate gyrus during theta oscillations in vivo. Neuroscience 57:555–564.

Stone SS, Teixeira CM, Zaslavsky K, Wheeler AL, Martinez-Canabal A, Wang AH, Sakaguchi M, Lozano AM, Frankland PW (2011) Functional convergence of developmentally and adult- generated granule cells in dentate gyrus circuits supporting hippocampus-dependent memory. Hippocampus 21:1348–1362.

Takao K, Miyakawa T (2006) Light/dark transition test for mice. J Vis Exp:104.

Teixeira CM, Rosen ZB, Suri D, Sun Q, Hersh M, Sargin D, Dincheva I, Morgan AA, Spivack S, Krok AC, Hirschfeld-Stoler T, Lambe EK, Siegelbaum SA, Ansorge MS (2018) Hippocampal 5-HT input regulates memory formation and schaffer collateral excitation. Neuron 98:992–1004 e1004.

Temmingh H, Stein DJ (2015) Anxiety in patients with schizophrenia: Epidemiology and management. CNS Drugs 29:819–832.

Vogel-Ciernia A, Wood MA (2014) Examining object location and object recognition memory in mice. Curr Protoc Neurosci 69:8 31 31–17.

Yagi S, Galea LAM (2019) Sex differences in hippocampal cognition and neurogenesis. Neuropsychopharmacology 44:200–213.

Yeh CY, Asrican B, Moss J, Quintanilla LJ, He T, Mao X, Casse F, Gebara E, Bao H, Lu W, Toni N, Song J (2018) Mossy cells control adult neural stem cell quiescence and maintenance through a dynamic balance between direct and indirect pathways. Neuron 99:493–510 e494.

Yuan Y, Wang H, Wei Z, Li W (2015) Impaired autophagy in hilar mossy cells of the dentate gyrus and its implication in schizophrenia. J Genet Genomics 42:1–8.

Zhou QG, Nemes AD, Lee D, Ro EJ, Zhang J, Nowacki AS, Dymecki SM, Najm IM, Suh H (2019) Chemogenetic silencing of hippocampal neurons suppresses epileptic neural circuits. J Clin Invest 129:310–323.

Zitman FM, Richter-Levin G (2013) Age and sex-dependent differences in activity, plasticity and response to stress in the dentate gyrus. Neuroscience 249:21–30.

